# Decomposing the role of alpha oscillations during brain maturation

**DOI:** 10.1101/2020.11.06.370882

**Authors:** Marius Tröndle, Tzvetan Popov, Sabine Dziemian, Nicolas Langer

## Abstract

Childhood and adolescence are critical stages of the human lifespan, in which fundamental neural reorganizational processes take place. A substantial body of literature investigated neurophysiological changes during brain maturation by focusing on the most dominant feature of the human EEG signal: the alpha oscillation. Ambiguous results were reported for the developmental trajectory of the power of the alpha oscillation. Simulations in this study show that conventional measures of alpha power are confounded by various factors and need to be decomposed into periodic and aperiodic components, which represent distinct underlying brain mechanisms. It is therefore unclear how each part of the signal relates to changes during brain maturation. Using multivariate Bayesian generalized linear mixed models, we examined aperiodic and periodic parameters of alpha activity in the largest openly available pediatric dataset (N=2529, age range 5-21 years) and replicated these findings in a preregistered analysis of an independent validation sample (N=369, age range 6–21yrs). First, the well documented age-related decrease in total alpha power was replicated. However, when controlling for the aperiodic signal component, our findings provide strong evidence for a reversed developmental trajectory of the periodic alpha power, whereas the aperiodic signal components, slope and offset, decreased. Consequently, earlier interpretations on age related changes of alpha power need to be fundamentally reconsidered, incorporating changes in the aperiodic signal. The interpretation of decreased total alpha power as elimination of active synapses rather links to decreases in the aperiodic intercept. Instead, additional analyses of diffusion tensor imaging data indicate that the maturational increase in periodic alpha power is related to increased thalamocortical connectivity. Functionally, our results suggest that increased thalamic control of cortical alpha power is linked to improved attentional performance during brain maturation.

## 1. Introduction

Childhood and adolescence are critical stages of the human lifespan, in which the brain undergoes various and complex micro- and macroscopic changes (Giedd et al., 1999; Lebel, Walker, Leemans, Phillips, & Beaulieu, 2008). The typical emergence of mental illnesses during childhood and adolescence (Kessler et al., 2005) further indicates fundamental maturational reorganization. It is therefore particularly important to understand these maturational changes in brain structure and function, which are accompanied by neurophysiological changes. A substantial body of literature has focused on investigating these neurophysiological changes with electroencephalography (EEG) (for reviews, see Anderson & Perone, 2018; Segalowitz, Santesso, & Jetha, 2010). Cognitive functions such as attention and memory, which undergo critical changes during maturation, have frequently been associated with EEG alpha activity (Foxe & Snyder, 2011; Klimesch, 1997; Klimesch, 2012; Niedermeyer, 1999). This lead to the notion that developmental changes in alpha activity reveals mechanisms of cortical manifestations of cognitive function. Indeed, numerous studies have investigated developmental changes in this alpha oscillation and reported evidence of an increase in the individual alpha frequency (IAF) at around 7 to 14 years of age (Cragg et al., 2011; Díaz de León, Harmony, Marosi, Becker, & Alvarez, 1988; Klimesch, 1999; Lindsley, 1939; Marcuse et al., 2008; Niedermeyer, 1999; Somsen, van’t Klooster, van der Molen, van Leeuwen, & Licht, 1997). However, the evidence is less clear about the amplitude of this alpha oscillation, termed alpha power. Absolute power was found to decrease with increasing age in some studies (Díaz de León et al., 1988; Gasser, Verleger, Bächer, & Sroka, 1988; Harmony et al., 1995; Lindsley, 1939; Whitford et al., 2007) but not in others (Clarke, Barry, McCarthy, & Selikowitz, 2001; Somsen et al., 1997). A potential confound is the utilization of fixed-frequency boundaries (e.g., 8–13 Hz), which neglects the slowing of the IAF during development. For instance, peak frequency in childhood is around 6 Hz but increases to 10 Hz in adolescents. Hence, age-related power decreases are underestimated when the slower alpha oscillation of younger children is not properly captured by predefined frequency limits, which will lead to lower power values. Consequently, individualized alpha frequency bands need to be extracted, which are centered on the individual IAF of each subject. In addition, thickening of the skull and other maturational processes that are unrelated to changes in neural activity manifest as changes in overall neural power. Thus, they pose a crucial confound in interpreting the relationship between alpha power and age. To overcome this latter limitation, studies have examined alpha power as relative to the overall power of the spectrum, relative alpha power, which has yielded more consistent results indicating an increase of alpha power with increasing age (Clarke et al., 2001; Cragg et al., 2011; Díaz de León et al., 1988; Harmony et al., 1995; John et al., 1980; Somsen et al., 1997). However, relative power measures of different frequency bands are by definition highly interdependent, as the frequency band of interest is normalized by the power of all the other frequency bands measured (Gómez et al., 2017; Somsen et al., 1997). For example, power changes in other frequency bands, such as the theta band, manifest as changes in relative alpha power, even though the true oscillatory alpha power may remain stable (see simulation analysis in supplement E and supplementary Figure 5A). Therefore, the extent to which these earlier works on alpha power and brain maturation are in effect confounded by changes in other frequency bands or by a slowing of the IAF remains unclear. To address this question, true alpha oscillatory power needs to be separated from other, non-oscillatory signal components.

Recent methodological developments have provided a means by which this separation can be achieved. These new approaches decompose measured power into periodic and aperiodic signal components (see Figure 4; Donoghue et al., 2020b; Hughes, Whitten, Caplan, & Dickson, 2012a; Wen & Liu, 2016). The aperiodic signal (i.e., “1/f signal”) is characterized by its intercept and slope, as its amplitude decreases with higher frequencies f. The aperiodic slope has been linked to the synchronicity of activity in the underlying neural population (Miller, Sorensen, Ojemann, & den Nijs, 2009; Usher, Stemmler, & Olami, 1995) and its balance of excitatory and inhibitory activity (Gao, Peterson, & Voytek, 2017), the aperiodic intercept has been linked to the general spiking activity (Voytek & Knight, 2015). Thus, the aperiodic signal contains important physiological information and needs to be considered during the analysis of spectral power, rather than measuring power relative to the absolute zero (e.g., Donoghue et al., 2020b). Applying this approach allows to extract an aperiodic-adjusted measure of alpha power, which is independent of oscillatory activity in other frequency bands and changes in overall power and aperiodic activity.

A recent study adopted this methodology and found no age-related decrease in the aperiodic-adjusted alpha power but did find age-related decreases in the aperiodic intercept and a flattening of the slope (He et al., 2019). Similar results were reported by Cellier et al. (2021). However, conventional measures of total or relative alpha power were either not reported (Cellier et al., 2021), or did not show any relation to age (He et al., 2019), hence comparisons and integration of these results with the large body of previous literature are not possible. Furthermore, due to comparatively small sample sizes in these studies, there is insufficient evidence for the reported significant association between age and aperiodic signal components and whether the aperiodic adjusted alpha-power truly remain stable in this period of life; or if too little statistical power was provided to detect changes in this newly emerging measure of alpha power. To overcome limitations of previous studies, we analyzed the currently largest openly available pediatric EEG data set (N = 2529), comprising children and adolescents aged between 5 and 21 years (Alexander et al., 2017) and validated the results in a second, preregistered analysis (https://osf.io/7uwy2) of a dataset consisting of 369 children aged between 6 and 21 years.

Based on animal studies investigating cortical and subcortical neural generators of alpha activity in adult animals (Bishop, 1936; Lopes da Silva, 1991; Steriade, Gloor, Llinás, Da Lopes Silva, & Mesulam, 1990), it is generally assumed that the thalamus and thalamocortical interactions strongly modulate cortical alpha activity. It can thus be hypothesized that changes in the alpha oscillation in the maturing brain are driven by structural changes of the thalamus and thalamocortical connectivity. However, to the best of our knowledge, no study investigated the relationship between anatomical maturation of the thalamus and its connectivity and alpha oscillatory power during childhood and adolescence. To close this gap, we further examined how thalamic structural changes relate to the observed changes in the different measures of alpha power. We employed magnet resonance images (MRI) to operationalize changes in the thalamus in terms of thalamic volume, and diffusion tensor imaging (DTI) to estimate thalamocortical connectivity by white matter integrity of the thalamic radiation. This was done in a subsample of the larger main dataset, for which magnet resonance images (MRI) and diffusion tensor imaging (DTI) data were additionally available.

Taken together, the present study aims to delineate the role of alpha oscillations during brain maturation by investigating conventional and newly emerging alpha oscillatory parameters, aperiodic signal components (see Table 2) and its underlying anatomical basis in a large sample of children and adolescents. Our first goal is to replicate previous literature that reported an age-related increase of the IAF, increased relative alpha power and decreased total alpha power during brain maturation. However, we hypothesize that the relation of alpha band power and brain maturation is no longer present when adjusting alpha power for the aperiodic signal component and for the age-related increase of the IAF. Additionally, based on previous literature we expect a decrease in the aperiodic intercept and a flattening of the aperiodic slope during brain maturation. Finally, as the aperiodic-adjusted individualized alpha power is unbiased by changes in other frequency bands and the aperiodic signal component, and therefore is likely to provide the most accurate reflection of the true oscillatory activity, we hypothesize that this measure shows a significant association with thalamic anatomical measures.

## 2. Results

### 2.1 Main analysis

The relation of age to the various parameters estimated in the Bayesian regression model are shown in Figure 1. A decrease in total individualized alpha power with increasing age was confirmed. Importantly, this relationship inverts when individualized alpha power is adjusted for the aperiodic signal, which then shows an age-related increase in power. In relative individualized alpha power, Figure 1 indicates positive age trend. An age-related increase of the IAF and a decrease of the aperiodic intercept and slope are also indicated (e.g., Figure 1 bottom row).

**Figure 1:**
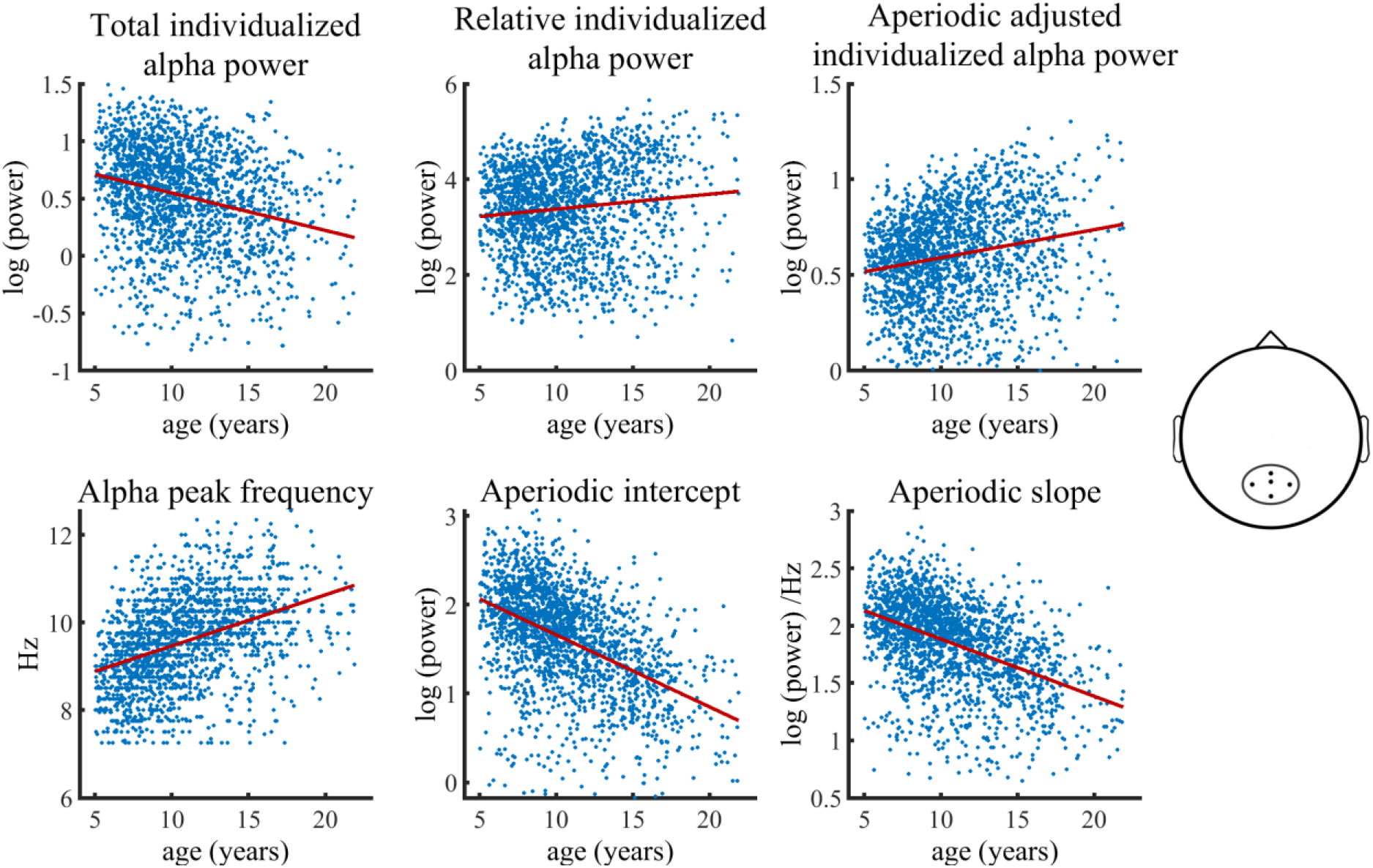
Visualization of data of the main HBN sample used in the Bayesian regression model. Solid lines represent fitted regression lines. The schematic head on the right indicates the location of the electrode cluster from which data was aggregated.

The statistical models further controlled for the heterogeneity of the full sample by adding a categorical diagnosis variable. An additional control analysis, in which we divided the ADHD diagnosis into two subdiagnoses of inattentive type and combined type, did not change any results (see supplementary Table 3). Most importantly, the control analysis, which included only the subsample with no diagnosis, showed results highly consistent with those of the full sample (see supplementary Figure 3 and supplementary Table 2).

The Bayesian regression model provided significant evidence for a reduction of total individualized alpha power during brain maturation (*b* = −0.31, *CI* = [−0.38, −0.24]). In contrast, a significant increase with increasing age was observed for aperiodic-adjusted alpha power (*b* = 0.23, *CI* = [0.16, 0.30]). Relative alpha power also showed a significant increase with increasing age (*b* = 0.14, *CI* = [0.06, 0.22]). The categorical diagnosis predictor did not show any significant effect on any of the outcomes. Females showed overall less power in total, relative, and aperiodic-adjusted individualized alpha and a lower aperiodic intercept in slope. Table 1 summarizes the results of the Bayesian regression model.

**Table 1.** Bayesian regression model results of full sample with categorical diagnosis predictor

| Outcome | age | gender | $\beta_{\text{predictor}}$ [CI] | | |
| --- | --- | --- | --- | --- | --- |
|  |  |  | diagnosis:<br>ADHD | diagnosis:<br>Other | age*gender |
| alpha peak frequency | 0.42 [0.34 - 0.49] | -0.08 [-0.15 - 0.02] | -0.09 [-0.19 - 0.01] | -0.07 [-0.17 - 0.04] | -0.04 [-0.15 - 0.08] |
| total individualized alpha power | -0.31 [-0.38 - 0.24] | -0.37 [-0.43 - 0.31] | 0.01 [-0.09 - 0.10] | 0.01 [-0.09 - 0.12] | 0.13 [0.01 - 0.25] |
| Relative individualized alpha power | 0.14 [0.06 - 0.22] | -0.35 [-0.41 - 0.28] | -0.01 [-0.11 - 0.10] | 0.01 [-0.10 - 0.12] | -0.05 [-0.17 - 0.07] |
| aperiodic-adjusted individualized alpha power | 0.23 [0.16 - 0.30] | -0.39 [-0.45 - 0.33] | -0.04 [-0.14 - 0.05] | -0.02 [-0.13 - 0.08] | -0.06 [-0.17 - 0.06] |
| aperiodic intercept | -0.54 [-0.61 - 0.48] | -0.37 [-0.42 - 0.32] | -0.02 [-0.10 - 0.07] | -0.02 [-0.10 - 0.08] | 0.08 [-0.02 - 0.19] |
| aperiodic slope | -0.44 [-0.51 - 0.38] | -0.39 [-0.44 - 0.33] | -0.04 [-0.12 - 0.05] | -0.03 [-0.12 - 0.06] | -0.05 [-0.15 - 0.06] |

*Note:* Credible Interval (CI) = 99.17%

The model also provided significant evidence of an age-related decrease in both the aperiodic slope and the aperiodic intercept. It was previously hypothesized that the diminished aperiodic intercept may merely be a result of a rotation of the power spectra due to a maturational flattening of the aperiodic slope (He et al., 2019). To test this hypothesis, we conducted an additional post hoc analysis in which median age split subsamples of the main HBN sample were bootstrapped 10 000 times, and the aperiodic signal of younger subjects was rotated by the corresponding age differences in aperiodic slope. The observed, rotated aperiodic intercepts were still larger than the actual aperiodic intercepts of corresponding older subjects in all of the 10 000 bootstrapped subsamples (mean age difference (older minus rotated younger) = −0.08, sd = 0.01, range = [−0.13 −0.04]). Hence, the observed decrease of the aperiodic intercept is larger than would be expected by shifts of the aperiodic slope (see supplement A for more details of the bootstrap analysis).

Analyses of canonically defined alpha power measures demonstrated a significant negative age effect on total canonical alpha power (*b* = −0.10, *CI*. [−0.17 −0.12]). Its effect size was significantly smaller than that of the age effect on aperiodic adjusted alpha power (*CI* is non-overlapping: [−0.38 −0.24]). Relative and aperiodic-adjusted canonical alpha power showed consistent significant age related increases. See supplementary Table 4 for detailed results on canonical alpha power

Figure 2 illustrates the age-related changes in the PSD during brain maturation. For visualization purposes, and in contrast to the statistical model, which used a continuous age variable, the grand averages for the youngest 20% (5.04–7.75 years) and oldest 20% (13.68–21.90 years) participants were plotted across the parieto-occipital electrodes. Figure 2A indicates an age-related decrease in alpha power in the total power spectrum, caused by lower total power values in older children in the lower frequency range of the alpha band (here ~7-10 Hz). Figure 2B visualizes the altered group differences when adjusting the power spectrum for the aperiodic signal. Here, an increase is observed in aperiodic-adjusted alpha power. The aperiodic signal shows a decreased intercept and flattened slope in age (Figure 2C).

**Figure 2:**
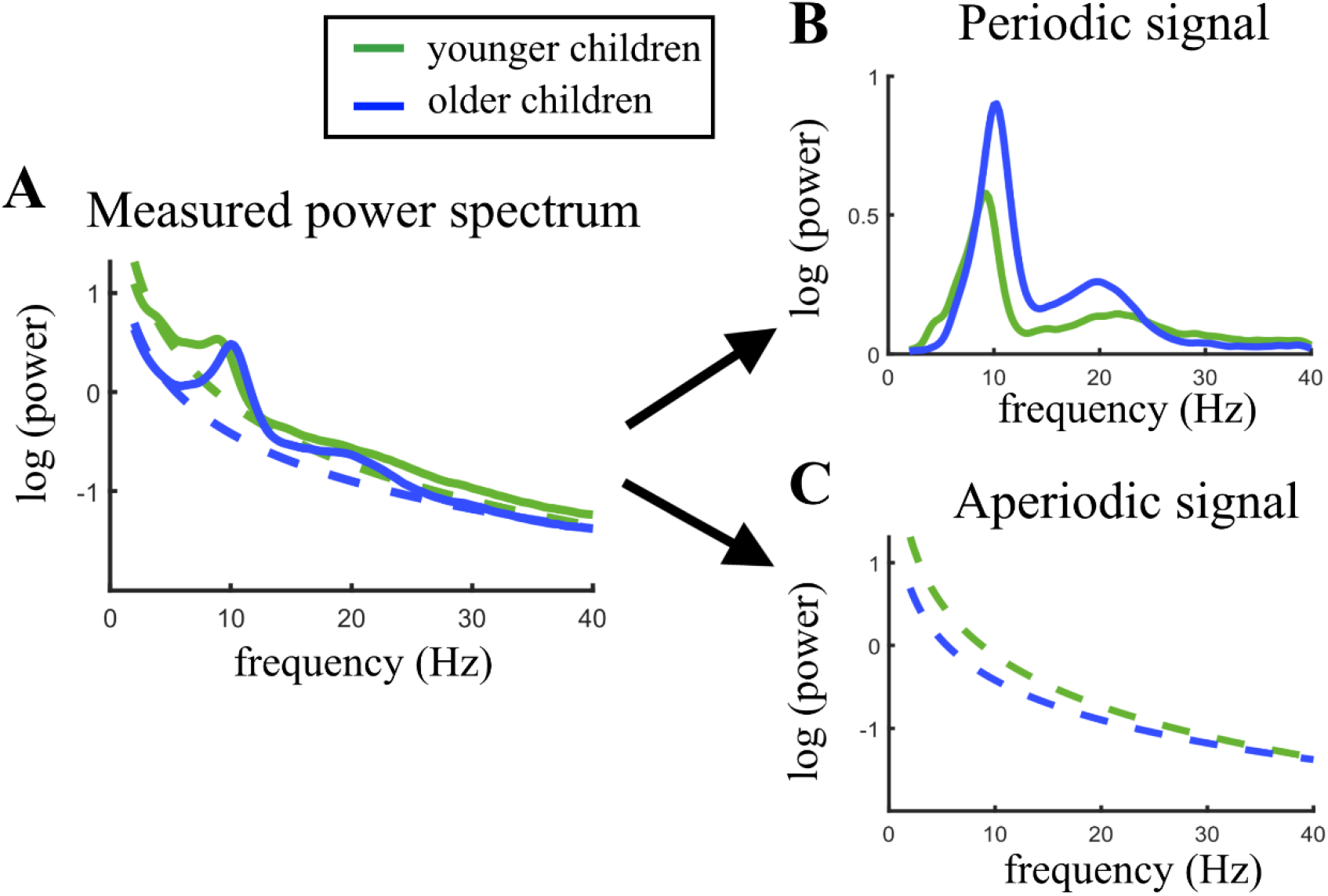
Visualization of age-related changes during brain maturation in (A) the measured power spectrum (i.e., total power spectrum), (B) the periodic (i.e., aperiodic-adjusted) power spectrum, and (C) the aperiodic signal. Younger children represent the 20% youngest children in the sample, older children the 20% oldest children and adolescents. This split of the sample was only done for visualization purposes and not used in any statistical analysis.

### 2.2 Validation Analyses

In the analyses of the validation sample, results were mostly consistent with those obtained from the main HBN sample. In terms of age effects, all results were replicated; however, the age-related decrease of relative individualized alpha power failed to reach significance. Here, neither gender nor the ADHD diagnosis showed any significant effect on the outcome measures. Table 2 summarizes the results of the statistical models using uninformative Cauchy priors.

**Table 2.** Bayesian regression model results of the full validation sample with categorical ADHD predictor

| Outcome | $\beta_{\text{predictor}}$ [CI] | | | |
| --- | --- | --- | --- | --- |
|  | age | gender | ADHD diagnosis | age*gender |
| alpha peak frequency | 0.34 [0.15 - 0.53] | -0.05 [-0.21 - 0.11] | -0.04 [-0.20 - 0.12] | 0.04 [-0.25 - 0.33] |
| total individualized alpha power | -0.44 [-0.61 - 0.27] | -0.05 [-0.19 - 0.11] | 0.04 [-0.11 - 0.19] | -0.07 [-0.33 - 0.30] |
| Relative individualized alpha power | 0.20 [0.00 - 0.39] | 0.05 [-0.12 - 0.22] | -0.02 [-0.18 - 0.15] | -0.10 [-0.40 - 0.20] |
| aperiodic-adjusted individualized alpha power | 0.33 [0.14 - 0.52] | -0.02 [-0.18 - 0.13] | -0.04 [-0.21 - 0.12] | -0.06 [-0.35 - 0.24] |
| aperiodic intercept | -0.76 [-0.88 - 0.65] | -0.10 [-0.19 - 0.00] | 0.00 [-0.10 - 0.09] | -0.07 [-0.25 - 0.11] |
| aperiodic slope | -0.60 [-0.77 - 0.45] | -0.08 [-0.21 - 0.05] | -0.06 [-0.19 - 0.06] | -0.13 [-0.36 - 0.10] |

*Note:* CI = 98.97% Credible Interval, gender variable is dummy coded: 1=female, 0=male.

Figure 3 visualizes age trajectories of the outcome measures in the full validation subsample.

**Figure 3:**
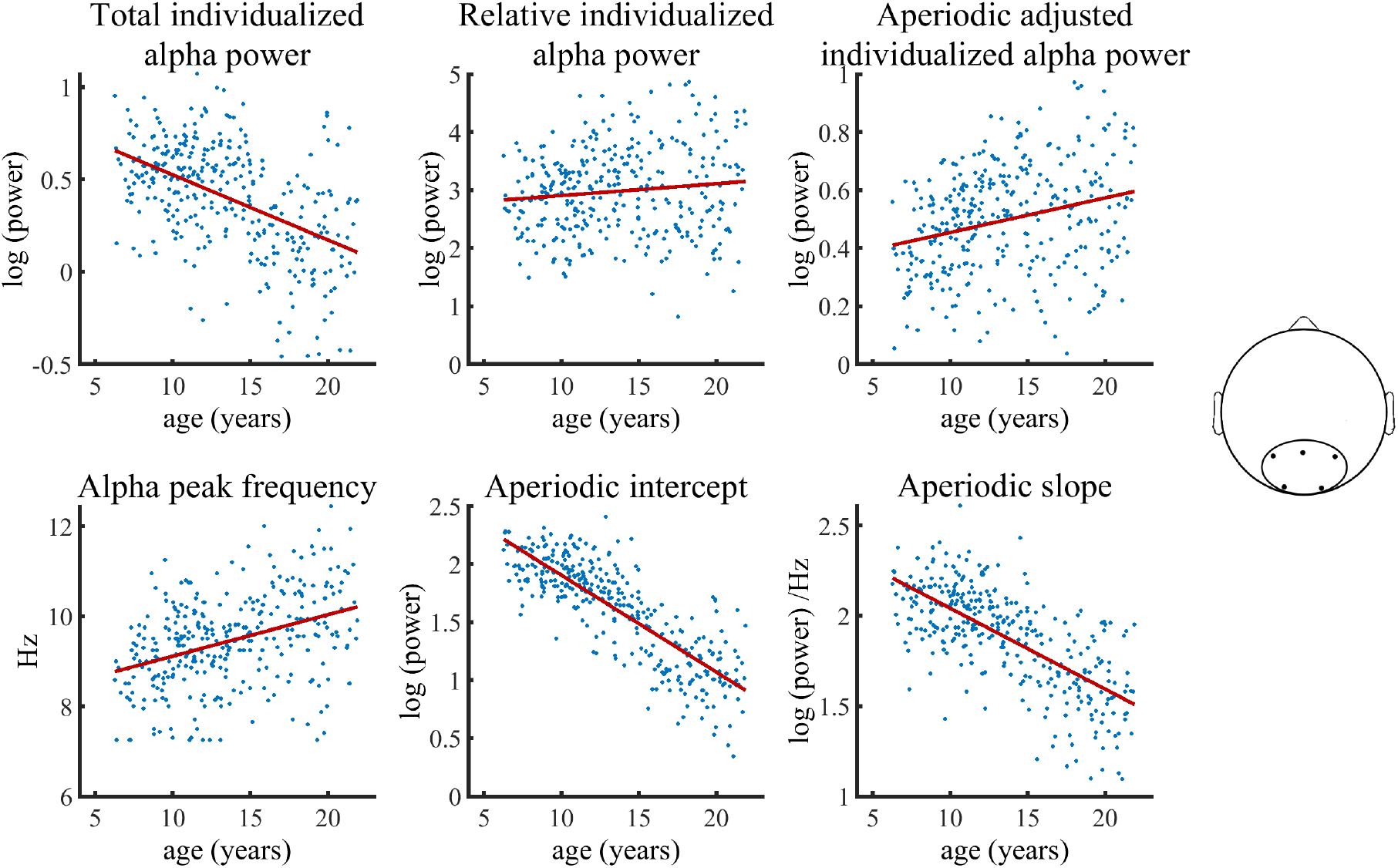
Visualization of data of the validation sample used in the Bayesian regression model. Solid lines represent fitted regression lines. The schematic head on the right indicates the location of the electrode cluster from which data was aggregated.

Importantly, the control analysis, which included only the healthy subsample, showed results highly consistent with those of the full sample (see supplementary Figure 4 and supplementary Table 5).

As an additional analysis, we applied Bayesian sequential updating to accumulate evidence across the different datasets. This procedure allows to increase statistical power and to produce more generalizable research outcome, which is less specific to each of the investigated samples. Therefore, informative priors extracted from the statistical results of the main sample were applied. Age effects on the six outcome variables in this analysis showed evidence consistent with the results obtained from the separate data sets. Gender showed significant effects on the three measures of alpha power and the aperiodic signal components here, although no gender effects were observable in the validation analyses using uninformative Cauchy priors. Because of the large difference between sample sizes (N_main_ = 1766, N_validation_ = 310), the extracted priors had a strong influence on the results; thus, it remains unclear how these gender effects generalize to other datasets. Clinical diagnosis of ADHD did not show any significant influence in the main and the validation analysis, however, in this analysis combining evidence across the two dataset, there was a significant negative relationship between ADHD diagnosis and aperiodic-adjusted alpha power, but not with aperiodic-adjusted or relative alpha power. See supplementary Table 6 for detailed results.

### 2.3 Relation of alpha power values to anatomical thalamic measures

Table 3 summarizes the results of the statistical models on the relation of thalamus volume and white matter integrity of the left and right thalamic radiation to total and aperiodic-adjusted alpha power. No relation was observed between thalamus volume and any outcome variable. Both the left and right thalamic radiation showed significant associations with aperiodic-adjusted individualized alpha power, which did not reach significance level with total individualized alpha power. Relative alpha power showed similar results as observed in aperiodic-adjusted alpha power.

**Table 3.** Summary table of the Bayesian regression model results on the influence of anatomical thalamic measures on total and aperiodic-adjusted alpha power

| Outcome | $\beta_{\text{predictor}}$ [CI] | | | | | |
| --- | --- | --- | --- | --- | --- | --- |
|  | left thalamic radiation | right thalamic radiation | thalamus volume | total intracranial volume | age | gender |
| total individualized alpha power | 0.07[-0.01 0.14] | 0.07 [0.00 0.15] | 0.00 [-0.11 0.12] | 0.03 [-0.07 0.12] | -0.26 [-0.34 - 0.18] | -0.32 [-0.41 - 0.23] |
| aperiodic-adjusted individualized alpha power | 0.15 [0.08 0.22] | 0.14 [0.07 0.21] | 0.03 [-0.08 0.14] | 0.02 [-0.08 0.11] | 0.17 [0.11 0.26] | -0.37 [-0.46 - 0.28] |

*Note:* CI = 97.37% Credible interval, gender variable is dummy coded: 1 = female, 0 = male. Credible intervals and $\beta$ estimates of the covariates (total intracranial volume, age, and gender) showed minor deviations across the three models (see 4.4.3, models 4, 5 & 6), due to the non-deterministic sampling in the Bayesian model estimation. Therefore, here they were averaged across the three models.

## 3. Discussion

This study investigated the role of alpha oscillations during brain maturation. Our results replicated the well-documented finding of an increasing IAF in this developmental phase of life from childhood to adolescence. The main aim of the study was to delineate the developmental trajectory of the power of the alpha oscillation by considering changes in the individual alpha frequency and aperiodic signal components in a large sample size. A significant decrease of total individualized alpha power was observed with increasing age across samples. However, when correcting for the aperiodic signal, the results changed considerably. The aperiodic-adjusted individualized alpha power increased significantly from childhood to adolescence. The aperiodic signal showed a decreased intercept with increasing age during brain maturation and a flattened slope. Results were largely consistent across the subsample of the HBN dataset without any given diagnosis, the full HBN dataset, and the validation analyses. The covariates of psychiatric diagnoses did not show any significant influences on oscillatory alpha or aperiodic signal parameters in the two datasets. Only in the Bayesian sequential updating analysis, combining both datasets, ADHD was associated with decreased aperiodic adjusted alpha power. Future research is needed to investigate possible relations between psychiatric diseases and periodic and aperiodic EEG signal components in more detail, which is beyond the scope of the present report. Gender effects on alpha power and aperiodic signal components were found in the HBN dataset; however, these effects did not generalize on the validation sample. Importantly, when relating alpha power measures to anatomical measures derived from DTI, only aperiodic-adjusted, but not total individualized alpha power showed a significant relation to the white matter integrity of the thalamic radiations.

### Age related changes in IAF and total canonical alpha power

The often-replicated increase of IAF from childhood to adolescence was also observed here. Higher IAF was previously linked to better sensorimotor abilities (Mierau et al., 2016) and increased memory performance (Klimesch, 1999). It was hypothesized that the mechanism underlying the increased memory performance is an increase in the speed of information processing, which was directly linked to the IAF by various studies investigating reaction time paradigms (Jin, O’Halloran, Plon, Sandman, & Potkin, 2006; e.g. Klimesch, Doppelmayr, Schimke, & Pachinger, 1996; Surwillo, 1961; Surwillo, 1961). This matches the findings of generally increased speed of information processing from childhood to adolescence (Kail, 2000). Thus, the increasing IAF observed during brain maturation may represent the neurophysiological correlate of increased speed of information processing. It has further been hypothesized that this is related to developmental increases in myelination and axon size (Cragg et al., 2011; Segalowitz et al., 2010).

In terms of alpha power analyses, the IAF increase induces a bias when alpha power is investigated with the canonical fixed-frequency bands, as was predominantly done in earlier investigations (Clarke et al., 2001; Díaz de León et al., 1988; Gasser et al., 1988; Harmony et al., 1995; Klimesch, 2012; Somsen et al., 1997; Whitford et al., 2007). Although some of these studies (Díaz de León et al., 1988; Gasser et al., 1988; Harmony et al., 1995; Whitford et al., 2007) found an age-related decrease in this total canonical alpha power measure, the present study only demonstrated a significant relationship between canonical alpha power and age in the large samples consisting of both healthy children and children diagnosed with ADHD or other psychiatric disorders. In contrast, the age effect on the alpha power adjusted for changes in IAF (i.e., total individualized alpha power) was significant across all datasets. It showed an effect size three times larger than that of the total canonical alpha power. Thus, the developmental decrease on total alpha power may be underestimated when using canonical frequency bands. This is because the slower alpha oscillation of younger children is not properly captured by predefined band limits and thus yields a lower alpha power value. Thus, we highly recommended refraining from using canonical fixed-frequency bands when investigating age-related changes in alpha power.

Even after accounting for the age-related changes in IAF, interpreting changes in total individualized alpha power still remains problematic. As noted earlier (e.g. Benninger, Matthis, & Scheffner, 1984; John et al., 1980; Matthis, Scheffner, Benninger, Lipinski, & Stolzis, 1980), a relative alpha power measure should be preferred over total alpha power, as the latter is more dependent on non-neurophysiological changes such as skull thickness and skin conductivity and is consequently less reliable (John et al., 1980).

### Relative vs. aperiodic-adjusted individualized alpha power

Relative individualized alpha power exhibited a positive relation to age in the full HBN sample, in contrast to the decrease in total individualized alpha power relative to the developmental trajectory. Thus, when overall changes in the power spectrum are taken into account, alpha power increased with increasing age. These results are in line with previous studies on relative alpha power (Clarke et al., 2001; Cragg et al., 2011; Díaz de León et al., 1988; Harmony et al., 1995; John et al., 1980; Somsen et al., 1997). However, normalizing alpha power by the power of all the other frequency bands measured poses problems for the interpretation of results. Post hoc simulations indicate that changes in power in other frequency bands (see supplementary Figure 5A) induce changes in relative alpha power even when true oscillatory alpha power is kept constant. Furthermore, changes in the aperiodic signal induce a confound in the relative alpha power measure (see supplementary Figure 5B). Consequently, changes in relative alpha power observed with increasing age need to be interpreted with caution, as changes in other frequency bands and in the aperiodic signal bias this finding. Our study confirmed an age-related decrease of the aperiodic intercept and a flattening of the aperiodic slope. Hence, relative alpha power measures are not conclusive indicators of a true age-related increase in alpha power. Instead, aperiodic-adjusted alpha power needs to be considered when analyzing developmental trajectories during brain maturation, as this measure of alpha power is independent of changes in both other frequency bands and the aperiodic signal.

### Age related increase in aperiodic-adjusted individualized alpha power

A significant increase of aperiodic-adjusted individualized alpha power with increasing age was observed across all samples. As discussed above, this measure of alpha power most likely reflects the true oscillatory changes in this phase of life, because it is independent of changes in the aperiodic signal and other frequency bands. Although other recent studies using smaller sample sizes and varying age ranges did not find an association of aperiodic-adjusted alpha power and brain maturation (Cellier et al., 2021; He et al., 2019), the present study provides strong evidence for an age-related increase across several large datasets.

Previous studies on age-related changes of alpha power during brain maturation speculated that decreased total oscillatory power may be due to synaptic pruning processes (e.g., Cragg et al., 2011) and thus reflect decreased spiking activity. The increase in aperiodic-adjusted alpha power contradicts these interpretations, decomposing the neural power spectra rather indicates that these processes relate to changes in the aperiodic signals intercept (see below). Another speculative hypothesis was that the observed developmental changes in the alpha band reflect structural changes in the thalamus and thalamocortical connectivity (Cragg et al., 2011; Whitford et al., 2007). However, this specific link was not formally investigated, but was based on early work on animal models, which explored adult cortical and subcortical neural generators of alpha activity. These models suggested that, in the adult animal brain, interacting thalamocortical loops may primarily be involved in the generation of the alpha rhythm (Bishop, 1936). Subsequent animal studies provided further evidence that the neural generators of alpha oscillations may be the occipital cortex, under strong guidance of the visual thalamus (Lopes da Silva, 1991; Lopes da Silva, van Lierop, Schrijer, & van Storm Leeuwen, 1973). So far, to the best of our knowledge, no studies were conducted to investigate if human maturational changes in the different measures of alpha power relate to anatomical changes in the thalamus and its connectivity. To close this missing link, we tested whether the resting-state alpha oscillatory power measures extracted here are related to anatomical measures of thalamic volume and thalamocortical connectivity, here measured by white matter integrity of the thalamic radiation. After age and total intracranial volume were controlled for, no association was found between thalamic volume and any of the alpha power measures. However, there was a significant positive relationship between white matter integrity of the thalamic radiation and aperiodic-adjusted individualized alpha power. Thus, the current study provides evidence that the increase in aperiodic-adjusted alpha power during brain maturation reflects increases in thalamocortical connectivity. The statistical models also indicated that this relation is not purely based on co-maturation of alpha power and structural connectivity, because significant relations were found between these measures after controlling for effects of age and the total intracranial volume. Total alpha power did not show any significant relation to these anatomical measures; hence, it is further indicated that this measure of alpha power is confounded by aperiodic signal components or non-neurophysiological changes during brain maturation. The finding that no significant relation between thalamus volume and alpha oscillatory power was found during brain maturation speculatively indicates that not the neural generators of alpha activity within the thalamus do change during brain maturation but that their interactions with cortical neural populations drive the changes in alpha oscillations.

Alterations in thalamocortical connectivity have been positively linked to attention and working memory performance in aging (Charlton, Barrick, Lawes, Markus, & Morris, 2010; Hughes et al., 2012b; Ystad et al., 2011) and infancy (Alcauter et al., 2014; Ball et al., 2015). Importantly, these cognitive functions have typically also been associated with the amplitude of alpha oscillation (Bazanova & Vernon, 2014; Foxe & Snyder, 2011; Klimesch, 1997; Klimesch, 1999; Klimesch, 2012) and are well known to improve during brain maturation. Furthermore, a highly influential simultaneous EEG and fMRI study (Laufs et al., 2003) hypothesized, based on correlations between cortical EEG and the blood oxygen level-dependent (BOLD) signal in frontoparietal regions, that adult resting state alpha power is linked to internally focused attention. These findings indicate that maturational increases of thalamic regulation of cortical alpha power may relate to improvements in attentional performance. As a matter of fact, post-hoc analyses (see supplement F) supported this hypothesis by providing evidence that oscillatory alpha power is linked to performance in a visual attention task. More specifically, the flanker task of the National Institutes of Health Toolbox Cognition Battery was employed as a measure of visual attention (Zelazo et al., 2013). While controlling for age, gender and handedness, aperiodic-adjusted individualized alpha power, but not total individualized alpha power, showed a significant positive relation to the attentional performance (see supplementary table 8). Taken together, contrary to findings based on total alpha power, aperiodic-adjusted individualized alpha power increased during brain maturation and seems likely to be related to increased thalamocortical connectivity. Functionally, this increase in alpha power is linked to improvements in attentional performance.

### Maturational changes in aperiodic signal components

The decreases in the aperiodic intercept and slope with increasing age are in line with previous observations in different age groups (Cellier et al., 2021; Donoghue, Dominguez, & Voytek, 2020a; He et al., 2019). Post hoc analyses showed that the diminished aperiodic intercept is not merely a result of a maturational flattening of the aperiodic slope, as hypothesized by He et al. (2019). One explanation for this decrease of the aperiodic intercept may lie within maturational changes that are unrelated to neural mechanisms. As previous studies on EEG changes during brain maturation have suggested (e.g.,Dustman, Shearer, & Emmerson, 1999), the thickening of the skull increases its resistance. This increased resistance could lead to the observed decrease in broadband EEG power, here reflected in a decrease in the aperiodic intercept. However, this maturational effect was also found in broadband power (Gómez et al., 2017) and the aperiodic intercept (He et al., 2019) in studies using magnetoencephalography (MEG), which is not affected by skull thickness.

Alternatively, as previous studies have shown that the aperiodic intercept is related to the overall spiking activity of the underlying neural population (Miller et al., 2009), the decrease observed in the aperiodic intercept may reflect a reduced parieto-occipital spiking activity during brain maturation. This observation may be related to the finding that as much as 40% of synapses in the striate cortex are eliminated during brain development (Huttenlocher & Courten, 1987). This is further supported by Whitford et al. (2007), who found a reduction of gray matter volume in parietal cortex co-occurring with a decrease in broadband EEG power during brain development, suggesting that this may reflect synaptic pruning processes. Yet, more research is needed to delineate possible mechanisms underlying this age-related decrease observed in the aperiodic intercept.

The flattening of the aperiodic signal may also be reflected in the commonly observed age-related decrease of power in low frequencies accompanied by an increase in power in higher frequencies in this age range (Cragg et al., 2011; Whitford et al., 2007). This phenomenon was speculated to be related to the elimination of synapses or changes in white matter structure (Segalowitz et al., 2010; Whitford et al., 2007), however, post hoc analyses did not show any relation of the aperiodic slope with white matter integrity of the thalamic radiation or global white matter integrity (see supplementary table 7). Yet, considering a shift of the aperiodic slope in the interpretation of this shift in power of frequency bands could provide additional insights into this little-understood result. Flattening of the aperiodic slope has been linked to changes in the excitation–inhibition ratio of the neural population (Gao et al., 2017), particularly an increase in local excitatory feedback. This shift in the excitation–inhibition ratio causes temporally decorrelated spikes and thus increases in neural noise (Voytek & Knight, 2015). This is supported by earlier studies relating a decreased aperiodic slope to more asynchronous activation patterns in neural populations (Miller et al., 2009; Usher et al., 1995). Thus, the flattened slope observed here might reflect increases in neural noise during brain development (McIntosh, 2010). This may seem contradictory, as aging research has linked an increase in neural noise to age-related cognitive decline from adulthood to old age (Voytek et al., 2015). However, increasing neural noise might rather have beneficial effects in the earlier processes of brain maturation. Reviewing maturational studies that used EEG and fMRI, McIntosh (2010) concluded that the maturing brain develops from a deterministic to a more stochastic system in which the increased neural noise leads to enhancement of functional network potential.

#### Limitations

A limitation of the present study is the composition of the samples investigated, as they contain a large proportion of children in whom psychiatric disorders were diagnosed. Yet, control analyses using only a healthy subsample show very similar results, additionally, categorical diagnoses were not significantly associated with the oscillatory and aperiodic signal components within both datasets.

### Conclusions

This study has demonstrated the relevance of taking the alpha peak frequency and aperiodic signal components into account when assessing age-related changes in spectral power during brain maturation. Our results indicate that there is significant variation of aperiodic activity during childhood and adolescence, which poses a confound to earlier work. Moreover, canonically defined frequency bands render age-related changes in IAF and power inseparable. Accounting for these confounding factors, and using the largest openly available pediatric sample, the present report demonstrates that the age effect on aperiodic-adjusted individualized alpha power shows the opposite direction as earlier assumed when investigating total alpha power. While previous recent studies did not find any relation between aperiodic-adjusted alpha power and age, the here applied robust statistical models provide strong evidence for an age related increase across large datasets. All results were confirmed in an independent validation dataset, indicating that aperiodic-adjusted individualized alpha power and the aperiodic signal intercept and slope are robust markers of the maturing brain. Previous interpretations indicated that maturational decreases in total alpha power may relate to the elimination of active synapses. Contrary to this, decomposition of the neural power spectrum reveals that these processes may rather relate to decreases of the aperiodic intercept. Instead, aperiodic-adjusted alpha power increases with increasing age, and likely reflects the development of thalamocortical connectivity. Functionally, these maturational changes may relate to increased attentional performance.

## 4. Materials and Methods

### 4.1. Datasets

For the main study, 2529 resting-state EEG datasets were obtained from the Human Brain Network (HBN) project (Alexander et al., 2017). The HBN project by the Child Mind Institute is an ongoing initiative that aims to generate a freely available biobank of multimodal datasets of children aged 5 to 21 years. These children undergo a variety of assessments. For the current study, we retrieved the Edinburgh handedness inventory (EHI, Oldfield, 1971), the Wechsler Intelligence Scale for Children-V (WISC-V, Wechsler, 2003) for children aged 6-17 or the Wechsler Adult Intelligence Scale (WAIS-IV, Wechsler, 2008) for adolescents older than 18 years, demographic data on age and gender, and clinical diagnoses. The clinical diagnoses are assessed by licensed clinicians who apply the DSM-5-based Schedule for Affective Disorders and Schizophrenia - Children’s version (KSADS) psychiatric interview (Kaufman et al., 1997). Additionally, when available, the corresponding MRI and DTI datasets were obtained (N = 786). Of the 2529 downloaded EEG datasets, 78 were not further processed because the EEG files were either corrupt or the recording length was not sufficient (i.e. the file size was smaller than 30 megabytes). Of the 2451 remaining EEG datasets, 178 could not be used because demographic data was missing. Applying the objective and reproducible exclusion criteria described below yielded a final sample size of 1766 subjects. See supplementary Figure 2 for a detailed flow chart of all exclusion criteria and the resulting sample size. Table 4 provides an overview of the final sample characteristics. Because the large proportion of children and adolescents with a diagnosed psychiatric disorder within the HBN sample might bias findings on general brain maturation, we conducted additional sensitivity analyses by subsampling this sample, including only subjects without any given diagnosis. This additional subsample consisted of 177 subjects.

For the preregistered validation study, a second dataset was employed which had previously been collected in a multicentric study. This dataset consisted of 369 healthy children and children with ADHD aged 6 to 21 years. In this sample, IQ was measured by CFT 1-R (Weiß, 2011) for children below the age of 9 years, CFT 20-R part I (Weiß, 2011) for adolescents between 9 and 16 years, and WMT-2 for adolescents older than 16 years (Forman, Poswanger, & Waldherr, 2006). Table 4 summarizes sample characteristics of the final sample (see supplementary Figure 1 for a visualization of the distribution of age and gender in the HBN and the validation sample).

**Table 4.**
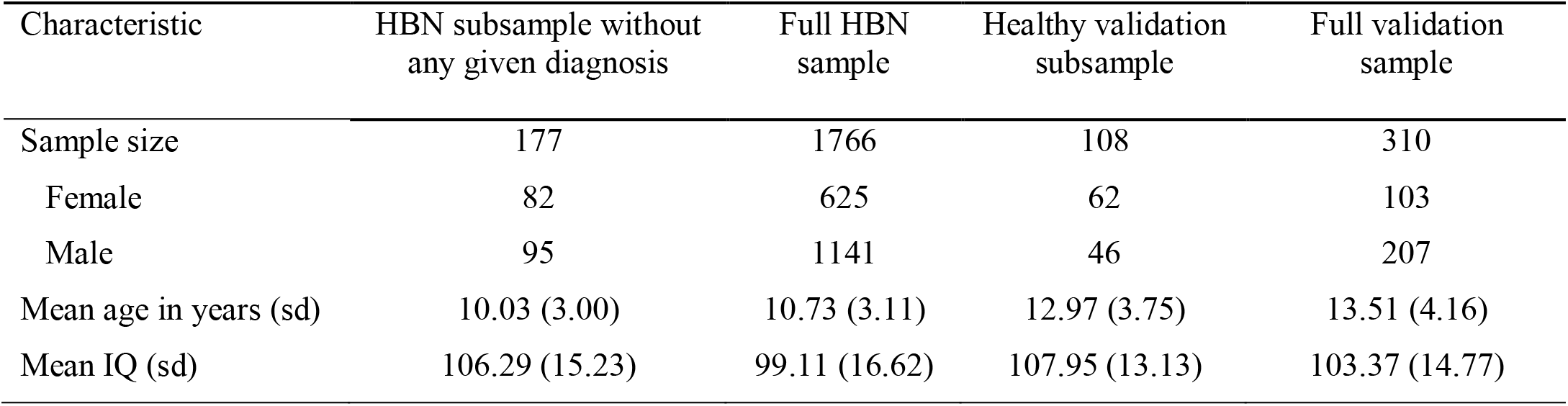
Characteristics of the final sample

### 4.2 Main Study

#### 4.2.1. Experimental setup and procedure

The participants of the HBN sample were comfortably seated in a chair in a sound-shielded room at a distance of 70 cm from a 17-inch CRT monitor (SONY Trinitron Multiscan G220, display dimensions 330 × 240 mm, resolution 800 × 600 pixels, vertical refresh rate of 100 Hz). The room was equipped with a chinrest to minimize head movements. Subjects were informed that EEG would be recorded while they rested with their eyes alternately open or closed. Instructions for the tasks were presented on the computer screen, and a research assistant answered questions from the participant from the adjacent control room through an intercom. Compliance with the task instructions was confirmed through a live video-feed to the control room. The task procedure was that participants rested with their eyes open for 20 s (a total of 1 min 40 sec), followed by 40 s (a total of 3 min 20 sec) with their eyes closed, repeated five times. Prerecorded verbal instructions automatically informed the participants when to open or close their eyes via loudspeakers. Participants were asked to maintain a fixed gaze on the fixation cross throughout EO blocks. The total duration of the EEG recording was 5 minutes. The alternating order of EO and EC was designed to avoid fatigue and maintain vigilance. The duration of EC blocks was twice as long as EO blocks because eyes-closed data is more robust and contains fewer artifacts. This protocol has been used in various previous studies (Langer et al., 2012; Langer, Bastian, Wirz, Oberauer, & Jäncke, 2013).

#### 4.2.2. Electroencephalography recording and preprocessing

The EEG was recorded at a sampling rate of 500 Hz using a high-density 128-channel EEG Geodesic Netamps system (Electrical Geodesics, Eugene, Oregon). The recording reference was at Cz, the vertex of the head, and impedances were kept below 40 kΩ. All analyses were performed using MATLAB 2018b (The MathWorks, Inc., Natick, Massachusetts, United States). EEG data was automatically preprocessed using the current version (2.4.3) of the MATLAB toolbox Automagic (Pedroni, Bahreini, & Langer, 2019). Our pipeline consisted of the following steps. First, bad channels were detected by the algorithms implemented in the clean_rawdata eeglab plugin (http://sccn.ucsd.edu/wiki/Plugin_list_process). A channel was defined as a bad electrode when data recorded by that electrode was correlated at less than 0.85 with an estimate based on other channels. Furthermore, a channel was defined as bad if it had more line noise relative to its signal than all other channels (four standard deviations). Finally, a channel was considered bad if it had a longer flat-line than 5 s. These bad channels were automatically removed and later interpolated using a spherical spline interpolation (EEGLAB function eeg_interp.m). The interpolation was later performed as a final step before the automatic quality assessment of the EEG files (see below). Next, data was filtered using a high-pass filter (−6dB cut off: 0.5 Hz). Line noise artifacts were removed by applying Zapline (Cheveigné, 2020), removing seven power line components. Subsequently, independent component analysis (ICA) was performed. Components reflecting artifactual activity were classified by the pretrained classifier ICLabel (Pion-Tonachini, Kreutz-Delgado, & Makeig, 2019). Components which were classified as any class of artifacts, including line noise, channel noise, muscle activity, eye activity, and heart artifacts, with a probability higher than 0.8 were removed from the data. Subsequently, residual bad channels were excluded if their standard deviation exceeded a threshold of 25 μV. Very high transient artifacts (> ±100 μV) were excluded from the calculation of the standard deviation of each channel. However, if this resulted in a significant loss of channel data (> 25%), the channel was removed from the data. After this, the pipeline automatically assessed the quality of the resulting EEG files based on four criteria: A data file was marked as bad-quality EEG and not included in the analysis if, first, the proportion of high-amplitude data points in the signals (>30 μV) was larger than 0.20; second, more than 20% of time points showed a variance larger than 15 microvolt across channels; third, 30% of the channels showed high variance (>15 μV); and fourth, the ratio of bad channels was higher than 0.3. Finally, 13 of 128 electrodes in the outermost circumference, attached to chin and neck, were excluded from further processing as they capture little brain activity and mainly record muscular activity. Additionally, 10 EOG electrodes were separated from the data and not used for further analysis, yielding a total of 105 EEG electrodes. Data was then referenced to the common average reference.

#### 4.2.3. Spectral analysis

Spectral analysis was performed on data from the concatenated five blocks of the eyes-closed condition. Only data from the eyes-closed condition was analyzed, because this data contains fewer artifacts and generally shows the strongest alpha oscillatory activity. The first and the last second of each eyes-closed block was discarded to exclude motor activity related to opening and closing the eyes and auditory activity due to the prompt from the speakers. The remaining data was concatenated, resulting in a total of 190 seconds of continuous EEG data. This data was again segmented into 2-second epochs, and each epoch containing large amplitude artifacts (> +90 μV, < −90 μV) was excluded from further processing. In 37 subjects, more than 50% of trials exceeded this threshold; thus, these subjects were not included in subsequent analyses (see supplementary Figure 2). For the remaining data, on average, 2.95% of trials were excluded by this criterion. Power spectral densities (PSDs) were then calculated using Welch’s Method (Welch, 1967) implemented in the EEGLab toolbox (Delorme & Makeig, 2004). Zero padding was applied to provide a frequency resolution of 0.25 Hz in the 2 s sliding time windows within Welch’s algorithm. Averaging the individual PSDs of each window resulted in a smoothed power spectrum that complies with the requirements of the specparam algorithm (Donoghue et al., 2020b) used subsequently (see 4.2.6). Additionally, PSDs were transformed to log scale to scale results equal to outputs from the specparam algorithm, which only operates in log power space. In the following, we describe the two approaches to extracting total alpha power and aperiodic-adjusted alpha power together with the aperiodic signal. See Table 5 for an overview of all extracted parameters.

**Table 5.** Overview of extracted parameters

| Parameter | Description |
| --- | --- |
| Individual alpha frequency (IAF) | Frequency at maximum power in search window 7–14Hz |
| Total canonical alpha power | Averaged log power in the fixed-frequency window [8Hz–13Hz], extracted from the total power spectrum |
| Total individualized alpha power | Averaged log power in window [- 4Hz to + 2Hz] relative to IAF, extracted from the total power spectrum |
| Relative canonical alpha power | Averaged power in the fixed-frequency window [8Hz–13Hz], divided by the average power of the full power spectrum (2–40Hz), extracted from the total power spectrum |
| Relative individualized alpha power | Averaged power in window [- 4Hz to + 2Hz] relative to IAF, divided by the average power of the full power spectrum (2–40Hz), extracted from the total power spectrum |
| Aperiodic-adjusted canonical alpha power | Canonical alpha power, extracted from the aperiodic-adjusted log power spectrum |
| Aperiodic-adjusted individual alpha power | Individualized alpha power, extracted from the aperiodic-adjusted log power spectrum |
| Aperiodic intercept | Intercept parameter of the aperiodic signal extracted by specparam |
| Aperiodic exponent | Exponent parameter (i.e., negative slope) of the aperiodic signal extracted by specparam |

*Note:* The term total power spectrum refers to the power spectrum as extracted from the data using Welch’s algorithm. The aperiodic-adjusted power spectrum results from a subtraction of the aperiodic signal from the total power spectrum.

#### 4.2.4 Computation of individual alpha peak frequency

The IAF was found by determining the frequency of maximum power between a lower and upper frequency limit. Following previous work, these frequencies limits were set to 7 and 14 Hz (Posthuma, Neale, Boomsma, & Geus, 2001; Smit, Wright, Hansell, Geffen, & Martin, 2006). If the peak was located at the border of the search range, no alpha peak was extracted for that subject, and the corresponding data was excluded from further analysis (67 subjects, see supplementary Figure 2).

#### 4.2.5 Extraction of total and relative alpha power

To replicate the results of previous published findings, this standard analysis approach included no adjustment for the aperiodic background signal. If an alpha peak was identified (see 4.2.4), individualized total alpha power was extracted by averaging log power in the defined window [−4 Hz to + 2 Hz] relative to the IAF (Klimesch, 1999). This individualized alpha power measure was chosen over a canonically defined alpha range, 8–13 Hz (Babiloni et al., 2020), as the shift of the IAF during maturation might introduce a bias when power is averaged within a fixed-frequency window. Canonical alpha band power was also extracted for supplementary analysis.

Relative individualized and relative canonical alpha power were calculated by dividing the corresponding total alpha power values by the average power of the full spectrum (2–40 Hz).

#### 4.2.6 Specparam algorithm and aperiodic-adjusted alpha power

The specparam algorithm (Donoghue et al., 2020b) parameterizes the neural power spectrum to separate neural oscillations from the aperiodic background signal. The algorithm estimates oscillatory peaks that are superimposed on the aperiodic background signal (see Figure 4) and are therefore measured relative to this rather than to the absolute zero. Consequently, the specparam algorithm parametrizes the PSD by iteratively fitting the aperiodic background curve (L) to the observed smoothed spectral signal, resulting in two parameters: the aperiodic intercept b and the aperiodic exponent *χ* (i.e. slope, the smaller *χ*, the flatter the spectrum).

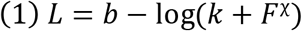

**Figure 4:**
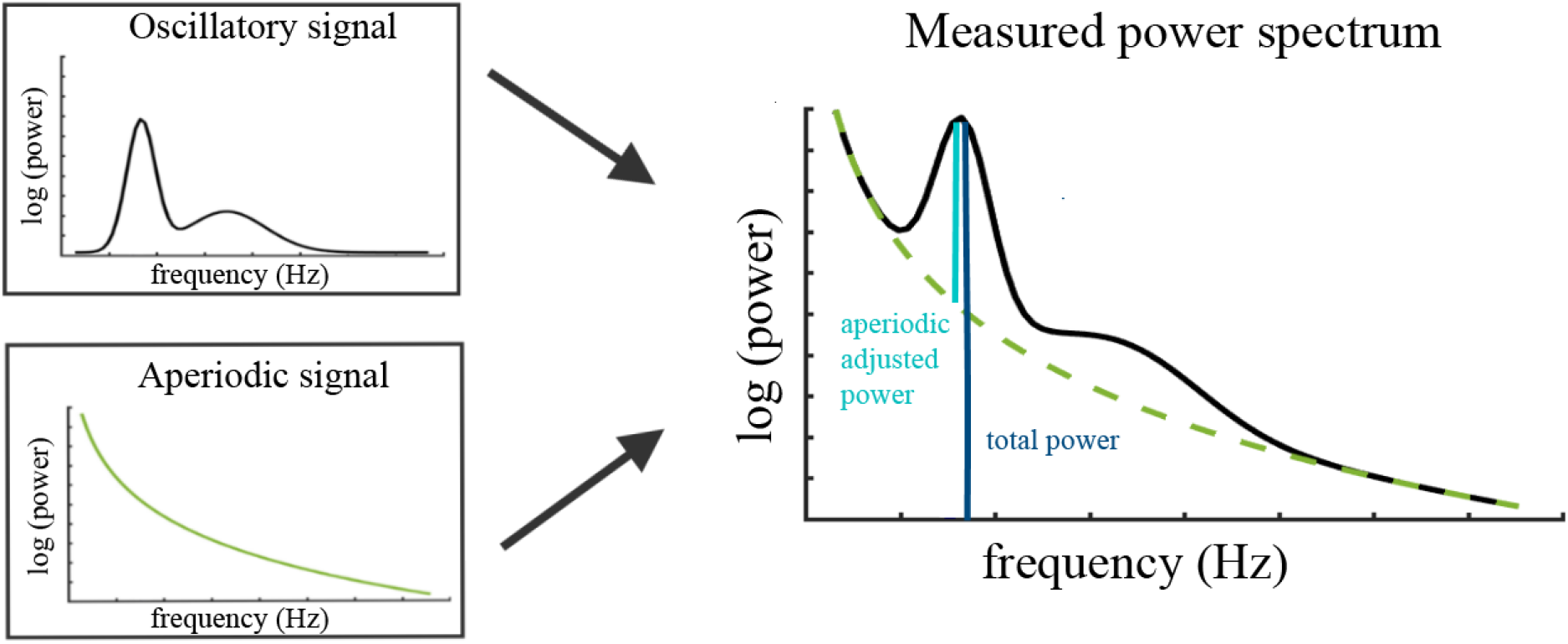
Illustration of the two components (left) superimposed in the measured neural power spectrum (right). The dark blue bar (right) indicates how total power is assessed relative to the absolute zero. The light blue bar represents aperiodic-adjusted power, which is assessed relative to the aperiodic signal.

In equation 1, F represents the vector of input frequencies and k the knee parameter, which is not further discussed here, as it is set to 0 in the proposed analysis: no bend of the aperiodic component is additionally modeled in the data, which is the default state of the specparam algorithm.

To extract oscillatory components, this aperiodic signal is subtracted from the power spectrum. Gaussians are fitted iteratively to the remaining signal and subsequently subtracted whenever data points exceed two standard deviations of the data. The Gaussians represent the true oscillatory components in the data; if data points are below the specified threshold, they are considered as noise. This results in a data-driven number of Gaussians, each parameterized by the frequency center, power relative to the aperiodic signal and the frequency bandwidth. The power spectrum is therefore modeled as defined by equation 2,

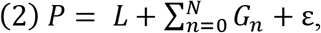

where *G*_n_ represents the *n*^th^ Gaussian and ε the noise not captured by the model. Note that this description of the algorithm is simplified; for a more detailed definition, see Donoghue et al. (2020b).

In the current study, the frequency range of 2 to 40 Hz of the power spectrum was passed to the algorithm because very low frequencies may lead to overfitting of noise as small bandwidth peaks. The current release (1.0.0) of the specparam toolbox from the github repository (https://github.com/fooof-tools/fooof) was used. The algorithm was used with these settings: peak width limits: [1, 8]; max number of peaks: infinite; minimum peak height: 0; peak threshold: 2 sd above mean; and aperiodic mode: fixed.

The resulting periodic signal P (“Oscillatory Signal” in Figure 4) was used to extract aperiodic-adjusted canonical alpha power (average across 8–13Hz) and aperiodic-adjusted individualized alpha power (average across a [−4 Hz to + 2 Hz] window centered on the IAF).

#### 4.2.7 Electrode cluster analysis

To test the hypothesis derived from literature review, an electrode-cluster-based analysis was performed. This cluster was based on data from the parietal and occipital electrodes, here referred to as the parieto-occipital cluster (see Figure 1). These electrodes were chosen because of the strong prominence of Oz and Pz electrodes in research on EEG alpha oscillations (Klimesch, 1999) and previous findings for age effects on alpha band power in these electrodes (Clarke et al., 2001; Cragg et al., 2011; Gasser et al., 1988). To account for individual anatomical differences, the following electrodes were selected to create a more robust cluster: E72 (POz), E75 (Oz), E62 (Pz), E67 (PO3), E77 (PO4). All parameters described above were extracted for each electrode and subsequently averaged within this cluster. Prior to statistical analyses, data was excluded from further processing if any extracted parameter exceeded a threshold of three standard deviations above or below the mean of the sample (excluded n=30, mean age =12.58, sd = 3.92). See supplementary Figure 2 for a detailed flow chart of all exclusion criteria and the resulting sample size.

#### 4.2.8 Statistical analysis

Bayesian generalized linear mixed models were formulated using the brms R package (Bürkner, 2017). Statistical models were separately fitted to the full sample and the subsampled dataset of subjects without any given diagnosis. For both samples, the predictor variable was age, and the covariates gender, EHQ, and site were added. In the full sample, a categorical variable for diagnosis was added (no diagnosis, ADHD diagnosis, other diagnosis) to account for the high prevalence of ADHD in this dataset (N_no diagnosis_: 177, N_ADHD diagnosis_: 1064, N_other diagnosis_: 525). However, the focus of this paper is on brain maturation; thus, this information is only included to control for possible confounding effects of psychiatric disorders.

To determine the best model for each analysis, multiple models were fitted with varying degrees of interaction terms and compared using the Watanabe Akaike Information Criteria (WAIC, Wantanabe, 2013). Additionally, expected log pointwise predictive density (ELDP, i.e. the expected predictive accuracy of the model) was calculated using the R function “loo_cmpare” (Vehtari, Gelman, & Gabry, 2017), yielding consistent results. For an overview of all models tested and the resulting WAIC and ELDP, see supplementary Table 1.

The final models were these:

For the subsample of subjects without any given diagnosis:

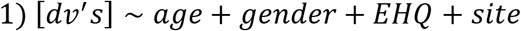

For the full sample:

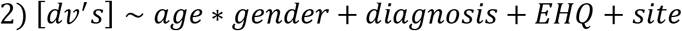

The multivariate models defined in equations 1 & 2 were fitted each to a set of dependent variables: Total individualized alpha power, aperiodic-adjusted individualized alpha power, relative individualized alpha power, aperiodic intercept, aperiodic slope and IAF. Additionally, for supplementary analyses, model two was also fitted to measures of canonical alpha power (i.e. total canonical alpha power, relative canonical alpha power and aperiodic adjusted canonical alpha power).

To correct for multiple comparisons, the significance level was adjusted. We assumed a high correlation between the total of 9 outcome variables (including the canonical alpha power measures, see supplementary Table 4), as many of the dependent variables represent various characteristics of the individual alpha oscillations. To account for this, we first calculated the effective number of tests of all dependent variables using Nyholt’s approach (Nyholt, 2004). Following this approach, the significance level (0.05) was then adjusted using the Šidák correction (Nyholt, 2004). Subsequently, the credible intervals (CIs) of the posterior distributions were calculated from the newly estimated levels of significance. The resulting significance level was 0.0083, yielding 99.17% credible intervals. We refrained from calculating Bayes factors for point estimates as evidence of the effect being zero or unequal to zero, as these Bayes factors, which are based on the Savage–Dickey ratio, depend strongly on the arbitrary choice of the prior distribution of each effect. Instead, we considered a model parameter significant if its 99.17% CI did not include zero. In line with Gelman’s recommendations (Gelman, Jakulin, Su, & Pittau, 2007), predictors and outcome variables of the Bayesian regression model were scaled as follows: Each numeric parameter (age, EHQ, IAF, total canonical alpha power, total individualized alpha power, relative canonical alpha power, relative individualized alpha power, aperiodic-adjusted canonical alpha power, aperiodic-adjusted individualized alpha power, aperiodic intercept, and aperiodic slope) was scaled to provide a mean of 0 and standard deviation 0.5. Uninformative Cauchy priors were used (mean = 0, sd = 2.5), as proposed by Gelman (Gelman et al., 2007).

### 4.3 Anatomical thalamic measures

Total and aperiodic-adjusted alpha power measures were further related to the thalamic volume and the fractional anisotropy (FA) of the thalamic radiation.

#### 4.3.1 MRI Data Acquisition

DTI and T1-weighted scans were acquired at three acquisition sites: Staten Island (SI), Rutgers University Brain Imaging Center (RUBIC), and CitiGroup Cornell Brain Imaging Center (CBIC). For the full scanning protocol and site-specific scanning parameters see Alexander et al. (2017) and supplement B.

#### 4.3.2 DTI Preprocessing

DTI data was processed using the FMRIB Software Library (FSL) version 6.0.4 (Jenkinson, Beckmann, Behrens, Woolrich, & Smith, 2012) following the “Diffusion parameter EstImation with Gibbs and NoisE Removal” (DESIGNER) pipeline (Ades-Aron et al., 2018). Detailed descriptions of all processing steps are available in supplement C and the entire preprocessing code is available at https://github.com/sdziem/DTIPreprocesingPipeline.

In brief, DTI data preprocessing included denoising (Veraart, Fieremans, & Novikov, 2016) followed by correction for Gibbs artefacts (Kellner, Dhital, Kiselev, & Reisert, 2016). Preprocessing continued with brain extraction with an FA threshold of 0.1 (Smith, 2002) and state-of-the-art correction for eddy current-induced distortions in-scanner head motion (Andersson et al., 2017; Andersson, Graham, Zsoldos, & Sotiropoulos, 2016; Andersson & Sotiropoulos, 2016). Next, outlier detection of MRI parameters and robust parameter estimation (Collier, Veraart, Jeurissen, Dekker, & Sijbers, 2015), tensor fitting, and extraction of diffusivity measures using weighted linear least squares estimation was applied (Fieremans, Jensen, & Helpern, 2011; Veraart et al., 2011; Veraart, Sijbers, Sunaert, Leemans, & Jeurissen, 2013).

Using automating fiber-tract quantification (AFQ, v.1.1) (Yeatman, Dougherty, Myall, Wandell, & Feldman, 2012), which implements a deterministic streamline tracking algorithm (Basser, Pajevic, Pierpaoli, Duda, & Aldroubi, 2000; Mori, Crain, Chacko, & van Zijl, 1999), we extracted objective and reliable FA values along the left and right thalamic radiation reflecting the tracts’ white matter integrity (Wassermann et al., 2011; Yeatman et al., 2011; Yeatman et al., 2012). In brief a co-registered T1-weighted scan was used to define anatomical regions of interest as seeds for consecutive tractography. Whole-brain tractography was computed using the following settings: FA threshold of 0.2, an FA mask threshold of 0.3, angle threshold of 35°. Next, the left and right thalamic radiations were segmented by distinct anatomical regions of interest as defined in a WM atlas (Wakana et al., 2007) in the co-registered T1-weighted scan. Fiber tract probability maps (Hua et al., 2008) were used to refine the tracts according to the likelihood of a fiber belonging to the tract. Fiber tract cleaning and outlier removal was performed using four standard deviations from the mean tract length and five standard deviations in distance from the tract core as removal criteria (Yeatman et al., 2012).

Each fiber belonging to the left and right thalamic radiation was sampled at 100 equidistant nodes (Yeatman et al., 2012). FA values were calculated for each segment along the tract as the sum of FA values of corresponding fibers weighted by the probability of the given fiber being part of the tract (Yeatman et al., 2012). This yields FA tract profiles for 100 equidistant segments for each participant’s left and right thalamic radiation. From these profiles, we calculated the tract-wise mean FA, which provides a more reliable quantification of the tract’s white matter integrity (Carlson et al., 2014; Luque Laguna et al., 2020).

Additionally, to ensure high imaging quality, each participant’s whole brain tractography underwent visual inspection for incomplete tractographies, gross artefacts, and misalignments of scans. Visual inspection was performed by a rater blind to the demographics of the participants. Only data rated as “good” was included in the final statistical analysis.

#### 4.3.3 T1-weighted Preprocessing

T1-weighted scans were preprocessed in parallel (Tange, 2011) with FreeSurfer (version 6.0.0) (http://surfer.nmr.mgh.harvard.edu/). Subcortical volumetric segmentation of the left and right thalamus was computed using the function “recon-all”. Details on the subcortical volumetric segmentation procedure have been reported previously (Fischl et al., 2002; Fischl et al., 2004; Han et al., 2006) and provided in supplement D. Briefly, processing included (1) correction for intensity non-uniformity, (2) Talairach transformation, (3) intensity normalization, (4) skull stripping, and (5) automated subcortical segmentation and labeling based on the default Gaussian Classifier Atlas (GCA) (Fischl et al., 2002; Fischl et al., 2004; Han et al., 2006). The validity of automated segmentation of the thalamus has been verified previously (Keller et al., 2012). Bilateral thalamic volumes (i.e., Left-Thalamus-Proper, Right-Thalamus-Proper) and total intracranial volume (i.e., EstimatedTotalIntraCranialVol) were extracted for subsequent analysis.

#### 4.3.4 Relation to alpha power

To investigate the relation between thalamic volume and the FA of the thalamic radiation, Bayesian regression models similar to those described in 4.2.8 were fitted, adding the thalamic volume and left and right thalamic radiation predictors while controlling for total intracranial volume, age & gender:

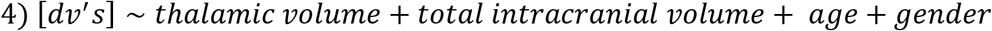

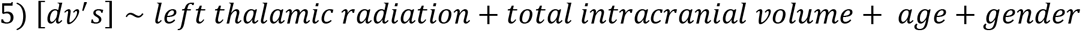

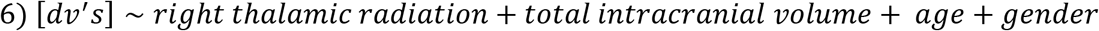

Due to collinearity issues, separate models were fitted for the left and right thalamic radiation. The dependent variables for Models 4, 5, and 6 were total individualized alpha power, aperiodic-adjusted individualized alpha power, and relative individualized alpha power. As described above, the numeric predictors and outcome variables were scaled to provide a mean of 0 and standard deviation 0.5. Uninformative Cauchy priors were used (mean = 0, sd = 2.5), as proposed by Gelman (Gelman et al., 2007). The models were fitted to the subset of the full sample, for which DTI for thalamic radiation and structural MRI for thalamic volume and total intracranial volume was available. This yielded a sample size of 592 subjects (mean age = 10.97, sd = 3.40, age range = 5.04 to 21.82, 209 female). To correct for multiplicity, the significance level was adjusted as described in section 4.2.8. For the three outcome variables, the resulting significance level was 0.0263, yielding 97.37% credible intervals.

### 4.4 Validation study

To validate the results from the analysis of the main dataset, the same analyses (sections 4.2.1–4.2.8) were applied to the second dataset of 369 subjects. Before the analysis pipeline was performed on this dataset, all analyses were preregistered in https://osf.io/7uwy2. In this dataset, eyes-closed resting-state EEG (mean length = 260s, sd = 28.4s) was recorded at a sampling rate of 500 Hz using a NeuroAmp ×23 with a PC-controlled 19-channel electroencephalographic system. Electrodes were placed according to the international 10–20 system using an electrode cap with tin electrodes (Electro-cap International Inc., Eaton, Ohio, USA). Data was referenced to the linked earlobes, and impedances were kept below 5 kΩ. Data was filtered between 0.5 and 50 Hz, and the same artifact correction as described in 4.2.2 was subsequently applied. The electrodes used for the parieto-occipital electrode cluster were Pz, P3, P4, O1, and O2. The same exclusion criteria as described above were applied. In this dataset, 34 subjects were excluded due to bad EEG data quality. Additionally, 11 subjects were excluded due to missing demographic data, 10 subjects had no detectable IAF, and 5 subjects were excluded due to outlier detection. This yielded a final sample size of 310 subjects (see Table 5 for an overview of the characteristics of the final sample). The parameters were scaled as described in 4.2.8, and the same uninformative Cauchy priors were used. An additional analysis combined evidence across the two datasets by extracting the posterior distributions for age and gender effects from the analyses of the HBN datasets and approximating them to the best-fitting distribution using the fitdistrplus R package (Delignette-Muller & Dutang, 2015). Subsequently, the statistical models were refitted using these extracted posteriors as priors for the analyses of the validation dataset.

## References

Ades-Aron, B., Veraart, J., Kochunov, P., McGuire, S., Sherman, P., Kellner, E., et al. (2018). Evaluation of the accuracy and precision of the diffusion parameter EStImation with Gibbs and NoisE removal pipeline. NeuroImage, 183, 532–543, from https://www.sciencedirect.com/science/article/pii/s1053811918306827.

Alcauter, S., Lin, W., Smith, J. K., Short, S. J., Goldman, B. D., Reznick, J. S., et al. (2014). Development of thalamocortical connectivity during infancy and its cognitive correlations. Journal of Neuroscience, 34(27), 9067–9075.

Alexander, L. M., Escalera, J., Ai, L., Andreotti, C., Febre, K., Mangone, A., et al. (2017). An open resource for transdiagnostic research in pediatric mental health and learning disorders. Scientific data, 4, 170181.

Anderson, A. J., & Perone, S. (2018). Developmental change in the resting state electroencephalogram: Insights into cognition and the brain. Brain and cognition, 126, 40–52.

Andersson, J. L. R., Graham, M. S., Drobnjak, I., Zhang, H., Filippini, N., & Bastiani, M. (2017). Towards a comprehensive framework for movement and distortion correction of diffusion MR images: Within volume movement. NeuroImage, 152, 450–466, from https://www.sciencedirect.com/science/article/pii/s1053811917301945.

Andersson, J. L. R., Graham, M. S., Zsoldos, E., & Sotiropoulos, S. N. (2016). Incorporating outlier detection and replacement into a non-parametric framework for movement and distortion correction of diffusion MR images. NeuroImage, 141, 556–572, from https://www.sciencedirect.com/science/article/pii/s1053811916303068.

Andersson, J. L. R., & Sotiropoulos, S. N. (2016). An integrated approach to correction for off-resonance effects and subject movement in diffusion MR imaging. NeuroImage, 125, 1063–1078, from https://www.sciencedirect.com/science/article/pii/s1053811915009209.

Babiloni, C., Barry, R. J., Başar, E., Blinowska, K. J., Cichocki, A., Drinkenburg, W. H. I. M., et al. (2020). International Federation of Clinical Neurophysiology (IFCN) - EEG research workgroup: Recommendations on frequency and topographic analysis of resting state EEG rhythms. Part 1: Applications in clinical research studies. Clinical neurophysiology: official journal of the International Federation of Clinical Neurophysiology, 131(1), 285–307.

Ball, G., Pazderova, L., Chew, A., Tusor, N., Merchant, N., Arichi, T., et al. (2015). Thalamocortical Connectivity Predicts Cognition in Children Born Preterm. Cerebral cortex (New York, N.Y.: 1991), 25(11), 4310–4318.

Basser, P. J., Pajevic, S., Pierpaoli, C., Duda, J., & Aldroubi, A. (2000). In vivo fiber tractography using DT-MRI data. Magnetic Resonance in Medicine, 44(4), 625–632.

Bazanova, O. M., & Vernon, D. (2014). Interpreting EEG alpha activity. Neuroscience and biobehavioral reviews, 44, 94–110, from https://www.sciencedirect.com/science/article/pii/s0149763413001279.

Benninger, C., Matthis, P., & Scheffner, D. (1984). EEG development of healthy boys and girls. Results of a longitudinal study. Electroencephalography and Clinical Neurophysiology, 57(1), 1–12.

Bishop, G. H. (1936). THE INTERPRETATION OF CORTICAL POTENTIALS. Cold Spring Harbor Symposia on Quantitative Biology, 4(0), 305–319, from http://symposium.cshlp.org/content/4/305.short.

Bürkner, P.-C. (2017). Advanced Bayesian Multilevel Modeling with the R Package brms, from http://arxiv.org/pdf/1705.11123v2.

Carlson, H. L., Laliberté, C., Brooks, B. L., Hodge, J., Kirton, A., Bello-Espinosa, L., et al. (2014). Reliability and variability of diffusion tensor imaging (DTI) tractography in pediatric epilepsy. Epilepsy & behavior: E&B, 37, 116–122, from https://www.sciencedirect.com/science/article/pii/s1525505014002261.

Cellier, D., Riddle, J., Petersen, I., & Hwang, K. (2021). The development of theta and alpha neural oscillations from ages 3 to 24 years. Developmental Cognitive Neuroscience, 50, 100969.

Charlton, R. A., Barrick, T. R., Lawes, I. N. C., Markus, H. S., & Morris, R. G. (2010). White matter pathways associated with working memory in normal aging. Cortex; a journal devoted to the study of the nervous system and behavior, 46(4), 474–489.

Cheveigné, A. de (2020). ZapLine: A simple and effective method to remove power line artifacts. NeuroImage, 207, 116356, from https://www.sciencedirect.com/science/article/pii/s1053811919309474.

Clarke, A. R., Barry, R. J., McCarthy, R., & Selikowitz, M. (2001). Age and sex effects in the EEG: development of the normal child. Clinical Neurophysiology, 112(5), 806–814, from http://www.sciencedirect.com/science/article/pii/S1388245701004886.

Collier, Q., Veraart, J., Jeurissen, B., Dekker, A. J. den, & Sijbers, J. (2015). Iterative reweighted linear least squares for accurate, fast, and robust estimation of diffusion magnetic resonance parameters. Magnetic Resonance in Medicine, 73(6), 2174–2184.

Cragg, L., Kovacevic, N., McIntosh, A. R., Poulsen, C., Martinu, K., Leonard, G., & Paus, T. (2011). Maturation of EEG power spectra in early adolescence: a longitudinal study. Developmental science, 14(5), 935–943, from https://pubmed.ncbi.nlm.nih.gov/21884309/.

Delignette-Muller, M. L., & Dutang, C. (2015). fitdistrplus: An R Package for Fitting Distributions. Journal of Statistical Software, 64(4).

Delorme, A., & Makeig, S. (2004). EEGLAB: an open source toolbox for analysis of single-trial EEG dynamics including independent component analysis. Journal of neuroscience methods, 134(1), 9–21.

Díaz de León, A. E., Harmony, T., Marosi, E., Becker, J., & Alvarez, A. (1988). Effect of different factors on EEG spectral parameters. The International journal of neuroscience, 43(1-2), 123–131.

Donoghue, T., Dominguez, J., & Voytek, B. (2020a). Electrophysiological Frequency Band Ratio Measures Conflate Periodic and Aperiodic Neural Activity. eNeuro.

Donoghue, T., Haller, M., Peterson, E., Varma, P., Sebastian, P., Gao, R., et al. (2020b). Parameterizing neural power spectra into periodic and aperiodic components. Nature neuroscience, in press.

Dustman, R. E., Shearer, D. E., & Emmerson, R. Y. (1999). Life-span changes in EEG spectral amplitude, amplitude variability and mean frequency. Clinical Neurophysiology, 110(8), 1399–1409, from https://www.sciencedirect.com/science/article/pii/s1388245799001029.

Fieremans, E., Jensen, J. H., & Helpern, J. A. (2011). White matter characterization with diffusional kurtosis imaging. NeuroImage, 58(1), 177–188, from https://www.sciencedirect.com/science/article/pii/s1053811911006148.

Fischl, B., Salat, D. H., Busa, E., Albert, M., Dieterich, M., Haselgrove, C., et al. (2002). Whole Brain Segmentation. Neuron, 33(3), 341–355, from https://www.sciencedirect.com/science/article/pii/s089662730200569x.

Fischl, B., Salat, D. H., van der Kouwe, A. J. W., Makris, N., Ségonne, F., Quinn, B. T., & Dale, A. M. (2004). Sequence-independent segmentation of magnetic resonance images. NeuroImage, 23 Suppl 1, S69–84, from https://www.sciencedirect.com/science/article/pii/s1053811904003817.

Forman, A. K., Poswanger, K., & Waldherr, K. (2006). Grundintelligenztest Skala 2 (CFT 20-R) mit Wortschatztest (WS) und Zahlenfolgentest (ZF).

Foxe, J. J., & Snyder, A. C. (2011). The Role of Alpha-Band Brain Oscillations as a Sensory Suppression Mechanism during Selective Attention. Frontiers in psychology, 2, 154.

Gao, R., Peterson, E. J., & Voytek, B. (2017). Inferring synaptic excitation/inhibition balance from field potentials. NeuroImage, 158, 70–78, from https://www.sciencedirect.com/science/article/pii/s1053811917305621.

Gasser, T., Verleger, R., Bächer, P., & Sroka, L. (1988). Development of the EEG of school-age children and adolescents. I. Analysis of band power. Electroencephalography and Clinical Neurophysiology, 69(2), 91–99.

Gelman, A., Jakulin, A., Su, Y.-S., & Pittau, M. G. (2007). A Default Prior Distribution for Logistic and Other Regression Models. SSRN Electronic Journal.

Giedd, J. N., Blumenthal, J., Jeffries, N. O., Castellanos, F. X., Liu, H., Zijdenbos, A., et al. (1999). Brain development during childhood and adolescence: a longitudinal MRI study. Nature neuroscience, 2(10), 861–863.

Gómez, C. M., Rodríguez-Martínez, E. I., Fernández, A., Maestú, F., Poza, J., & Gómez, C. (2017). Absolute Power Spectral Density Changes in the Magnetoencephalographic Activity During the Transition from Childhood to Adulthood. Brain Topography, 30(1), 87–97.

Han, X., Jovicich, J., Salat, D., van der Kouwe, A., Quinn, B., Czanner, S., et al. (2006). Reliability of MRI-derived measurements of human cerebral cortical thickness: the effects of field strength, scanner upgrade and manufacturer. NeuroImage, 32(1), 180–194, from https://www.sciencedirect.com/science/article/pii/s1053811906001601.

Harmony, T., Marosi, E., Becker, J., Rodríguez, M., Reyes, A., Fernández, T., et al. (1995). Longitudinal quantitative EEG study of children with different performances on a reading-writing test. Electroencephalography and Clinical Neurophysiology, 95(6), 426–433.

He, W., Donoghue, T., Sowman, P. F., Seymour, R. A., Brock, J., Crain, S., et al. (2019). Co-Increasing Neuronal Noise and Beta Power in the Developing Brain.

Hua, K., Zhang, J., Wakana, S., Jiang, H., Li, X., Reich, D. S., et al. (2008). Tract probability maps in stereotaxic spaces: analyses of white matter anatomy and tract-specific quantification. NeuroImage, 39(1), 336–347, from https://www.sciencedirect.com/science/article/pii/s105381190700688x.

Hughes, A. M., Whitten, T. A., Caplan, J. B., & Dickson, C. T. (2012a). BOSC: a better oscillation detection method, extracts both sustained and transient rhythms from rat hippocampal recordings. Hippocampus, 22(6), 1417–1428.

Hughes, E. J., Bond, J., Svrckova, P., Makropoulos, A., Ball, G., Sharp, D. J., et al. (2012b). Regional changes in thalamic shape and volume with increasing age. NeuroImage, 63(3), 1134–1142.

Huttenlocher, P. R., & Courten, C. de (1987). The development of synapses in striate cortex of man. Human neurobiology, 6(1), 1–9.

Jenkinson, M., Beckmann, C. F., Behrens, T. E. J., Woolrich, M. W., & Smith, S. M. (2012). FSL. NeuroImage, 62(2), 782–790, from https://www.sciencedirect.com/science/article/pii/s1053811911010603.

Jin, Y., O’Halloran, J. P., Plon, L., Sandman, C. A., & Potkin, S. G. (2006). Alpha EEG predicts visual reaction time. The International journal of neuroscience, 116(9), 1035–1044.

John, E. R., Ahn, H., Prichep, L., Trepetin, M., Brown, D., & Kaye, H. (1980). Developmental equations for the electroencephalogram. Science (New York, N.Y.), 210(4475), 1255–1258, from https://pubmed.ncbi.nlm.nih.gov/7434026/.

Kail, R. (2000). Speed of Information Processing. Journal of School Psychology, 38(1), 51–61.

Kaufman, J., Birmaher, B., Brent, D., Rao, U., Flynn, C., Moreci, P., et al. (1997). Schedule for Affective Disorders and Schizophrenia for School-Age Children-Present and Lifetime Version (K-SADS-PL): initial reliability and validity data. Journal of the American Academy of Child & Adolescent Psychiatry, 36(7), 980–988.

Keller, S. S., Gerdes, J. S., Mohammadi, S., Kellinghaus, C., Kugel, H., Deppe, K., et al. (2012). Volume estimation of the thalamus using freesurfer and stereology: consistency between methods. Neuroinformatics, 10(4), 341–350, from https://link.springer.com/article/10.1007/s12021-012-9147-0?error=cookies_not_supported&error=cookies_not_supported&code=8eb3577d-7908-479a-b6d3-a7df89471031&code=2304e0c3-d704-4d47-9699-ad1172a815ec.

Kellner, E., Dhital, B., Kiselev, V. G., & Reisert, M. (2016). Gibbs-ringing artifact removal based on local subvoxel-shifts. Magnetic Resonance in Medicine, 76(5), 1574–1581.

Kessler, R. C., Berglund, P., Demler, O., Jin, R., Merikangas, K. R., & Walters, E. E. (2005). Lifetime prevalence and age-of-onset distributions of DSM-IV disorders in the National Comorbidity Survey Replication. Archives of general psychiatry, 62(6), 593–602.

Klimesch, W., Doppelmayr, M., Schimke, H., & Pachinger, T. (1996). Alpha Frequency, Reaction Time, and the Speed of Processing Information. Journal of Clinical Neurophysiology, 13(6), 511, from https://journals.lww.com/clinicalneurophys/fulltext/1996/11000/alpha_frequency,_reaction_time,_and_the_speed_of.6.aspx.

Klimesch, W. (1997). EEG-alpha rhythms and memory processes. International Journal of Psychophysiology, 26(1-3), 319–340.

Klimesch, W. (1999). EEG alpha and theta oscillations reflect cognitive and memory performance: a review and analysis. Brain Research Reviews, 29(2-3), 169–195.

Klimesch, W. (2012). α-band oscillations, attention, and controlled access to stored information. Trends in cognitive sciences, 16(12), 606–617, from https://www.sciencedirect.com/science/article/pii/s1364661312002434.

Langer, N., Bastian, C. C. von, Wirz, H., Oberauer, K., & Jäncke, L. (2013). The effects of working memory training on functional brain network efficiency. Cortex; a journal devoted to the study of the nervous system and behavior, 49(9), 2424–2438, from http://www.sciencedirect.com/science/article/pii/S0010945213000117.

Langer, N., Pedroni, A., Gianotti, L. R. R., Hänggi, J., Knoch, D., & Jäncke, L. (2012). Functional brain network efficiency predicts intelligence. Human brain mapping, 33(6), 1393–1406.

Laufs, H., Kleinschmidt, A., Beyerle, A., Eger, E., Salek-Haddadi, A., Preibisch, C., & Krakow, K. (2003). EEG-correlated fMRI of human alpha activity. NeuroImage, 19(4), 1463–1476.

Lebel, C., Walker, L., Leemans, A., Phillips, L., & Beaulieu, C. (2008). Microstructural maturation of the human brain from childhood to adulthood. NeuroImage, 40(3), 1044–1055.

Lindsley, D. B. (1939). A Longitudinal Study of the Occipital Alpha Rhythm in Normal Children: Frequency and Amplitude Standards. The Pedagogical Seminary and Journal of Genetic Psychology, 55(1), 197–213.

Lopes da Silva, F., van Lierop, T., Schrijer, C., & van Storm Leeuwen, W. (1973). Organization of thalamic and cortical alpha rhythms: Spectra and coherences. Electroencephalography and Clinical Neurophysiology, 35(6), 627–639.

Lopes da Silva, F. (1991). Neural mechanisms underlying brain waves: from neural membranes to networks. Electroencephalography and Clinical Neurophysiology, 79(2), 81–93.

Luque Laguna, P. A., Combes, A. J. E., Streffer, J., Einstein, S., Timmers, M., Williams, S. C. R., & Dell’Acqua, F. (2020). Reproducibility, reliability and variability of FA and MD in the older healthy population: A test-retest multiparametric analysis. NeuroImage. Clinical, 26, 102168.

Marcuse, L. V., Schneider, M., Mortati, K. A., Donnelly, K. M., Arnedo, V., & Grant, A. C. (2008). Quantitative analysis of the EEG posterior-dominant rhythm in healthy adolescents. Clinical Neurophysiology, 119(8), 1778–1781.

Matthis, P., Scheffner, D., Benninger, C., Lipinski, C., & Stolzis, L. (1980). Changes in the background activity of the electroencephalogram according to age. Electroencephalography and Clinical Neurophysiology, 49(5-6), 626–635.

McIntosh, A. R. (2010). The Development of a Noisy Brain. Archives Italiennes de Biologie. (148), 223–337.

Mierau, A., Felsch, M., Hülsdünker, T., Mierau, J., Bullermann, P., Weiß, B., & Strüder, H. K. (2016). The interrelation between sensorimotor abilities, cognitive performance and individual EEG alpha peak frequency in young children. Clinical neurophysiology: official journal of the International Federation of Clinical Neurophysiology, 127(1), 270–276.

Miller, K. J., Sorensen, L. B., Ojemann, J. G., & den Nijs, M. (2009). Power-law scaling in the brain surface electric potential. PLoS computational biology, 5(12), e1000609.

Mori, S., Crain, B. J., Chacko, V. P., & van Zijl, P. C. M. (1999). Three-dimensional tracking of axonal projections in the brain by magnetic resonance imaging. Annals of Neurology, 45(2), 265–269.

Niedermeyer, E. (1999). The normal EEG of the waking adult. Electroencephalography: Basic principles, clinical applications, and related fields, 167, 155–164.

Nyholt, D. R. (2004). A simple correction for multiple testing for single-nucleotide polymorphisms in linkage disequilibrium with each other. American journal of human genetics, 74(4), 765–769.

Oldfield, R. C. (1971). The assessment and analysis of handedness: The Edinburgh inventory. Neuropsychologia, 9(1), 97–113, from https://www.sciencedirect.com/science/article/pii/0028393271900674.

Pedroni, A., Bahreini, A., & Langer, N. (2019). Automagic: Standardized preprocessing of big EEG data. NeuroImage, 460–473.

Pion-Tonachini, L., Kreutz-Delgado, K., & Makeig, S. (2019). ICLabel: An automated electroencephalographic independent component classifier, dataset, and website. NeuroImage, 198, 181–197, from https://www.sciencedirect.com/science/article/pii/s1053811919304185.

Posthuma, D., Neale, M. C., Boomsma, D. I., & Geus, E. J. de (2001). Are smarter brains running faster? Heritability of alpha peak frequency, IQ, and their interrelation. Behavior genetics, 31(6), 567–579.

Segalowitz, S. J., Santesso, D. L., & Jetha, M. K. (2010). Electrophysiological changes during adolescence: a review. Brain and cognition, 72(1), 86–100.

Smit, C. M., Wright, M. J., Hansell, N. K., Geffen, G. M., & Martin, N. G. (2006). Genetic variation of individual alpha frequency (IAF) and alpha power in a large adolescent twin sample. International journal of psychophysiology: official journal of the International Organization of Psychophysiology, 61(2), 235–243, from http://www.sciencedirect.com/science/article/pii/S0167876005002618.

Smith, S. M. (2002). Fast robust automated brain extraction. Human brain mapping, 17(3), 143–155.

Somsen, R. J., van’t Klooster, B. J., van der Molen, M. W., van Leeuwen, H. M., & Licht, R. (1997). Growth spurts in brain maturation during middle childhood as indexed by EEG power spectra. Biological Psychology, 44(3), 187–209.

Steriade, M., Gloor, P., Llinás, R. R., Da Lopes Silva, F. H., & Mesulam, M.-M. (1990). Basic mechanisms of cerebral rhythmic activities. Electroencephalography and Clinical Neurophysiology, 76(6), 481–508.

Surwillo, W. (1961). Frequency of the ‘Alpha’ Rhythm, Reaction Time and Age. Nature, 191(4790), 823–824, from https://www.nature.com/articles/191823a0.

Tange, O. (2011). Gnu parallel-the command-line power tool. The USENIX Magazine. (36 (1)), 42–47.

Usher, Stemmler, & Olami (1995). Dynamic pattern formation leads to 1/f noise in neural populations. Physical review letters, 74(2), 326–329.

Vehtari, A., Gelman, A., & Gabry, J. (2017). Practical Bayesian model evaluation using leave-one-out cross-validation and WAIC. Statistics and Computing, 27(5), 1413–1432.

Veraart, J., Fieremans, E., & Novikov, D. S. (2016). Diffusion MRI noise mapping using random matrix theory. Magnetic Resonance in Medicine, 76(5), 1582–1593.

Veraart, J., Poot, D. H. J., van Hecke, W., Blockx, I., van der Linden, A., Verhoye, M., & Sijbers, J. (2011). More accurate estimation of diffusion tensor parameters using diffusion Kurtosis imaging. Magnetic Resonance in Medicine, 65(1), 138–145.

Veraart, J., Sijbers, J., Sunaert, S., Leemans, A., & Jeurissen, B. (2013). Weighted linear least squares estimation of diffusion MRI parameters: strengths, limitations, and pitfalls. NeuroImage, 81, 335–346, from https://www.sciencedirect.com/science/article/pii/s1053811913005223.

Voytek, B., & Knight, R. T. (2015). Dynamic network communication as a unifying neural basis for cognition, development, aging, and disease. Biological psychiatry, 77(12), 1089–1097.

Voytek, B., Kramer, M. A., Case, J., Lepage, K. Q., Tempesta, Z. R., Knight, R. T., & Gazzaley, A. (2015). Age-Related Changes in 1/f Neural Electrophysiological Noise. Journal of Neuroscience, 35(38), 13257–13265.

Wakana, S., Caprihan, A., Panzenboeck, M. M., Fallon, J. H., Perry, M., Gollub, R. L., et al. (2007). Reproducibility of quantitative tractography methods applied to cerebral white matter. NeuroImage, 36(3), 630–644, from https://www.sciencedirect.com/science/article/pii/s1053811907001383.

Wantanabe, S. (2013). A widely applicable Bayesian information criterion. Journal of Machine Learning Research. (14), 867–897, from https://www.jmlr.org/papers/volume14/watanabe13a/watanabe13a.pdf.

Wassermann, D., Rathi, Y., Bouix, S., Kubicki, M., Kikinis, R., Shenton, M., & Westin, C.-F. (2011). White Matter Bundle Registration and Population Analysis Based on Gaussian Processes. In (pp. 320–332). Springer, Berlin, Heidelberg.

Wechsler, D. (2003). Wechsler intelligence scale for children--Fourth Edition (WISC-IV). The Psychological Corporation.

Wechsler, D. (2008). Wechsler Adult Intelligence Scale--Fourth edition (WAIS-IV), Australian and New Zealand Language Adaptation. NCS Pearson Inc.

Weiß, R. H. (2011). Wiener Matrizen-Test 2: Ein Rasch-skaldierter sprachfreier Kurztest zu Erfassung der Intelligenz.

Welch, P. (1967). The use of fast Fourier transform for the estimation of power spectra: A method based on time averaging over short, modified periodograms. IEEE Transactions on Audio and Electroacoustics, 15(2), 70–73.

Wen, H., & Liu, Z. (2016). Separating Fractal and Oscillatory Components in the Power Spectrum of Neurophysiological Signal. Brain Topography, 29(1), 13–26.

Whitford, T. J., Rennie, C. J., Grieve, S. M., Clark, C. R., Gordon, E., & Williams, L. M. (2007). Brain maturation in adolescence: concurrent changes in neuroanatomy and neurophysiology. Human brain mapping, 28(3), 228–237.

Yeatman, J. D., Dougherty, R. F., Myall, N. J., Wandell, B. A., & Feldman, H. M. (2012). Tract profiles of white matter properties: automating fiber-tract quantification. PLOS ONE, 7(11), e49790, from https://jourals.plos.org/plosone/article?id=10.1371/joural.pone.0049790.

Yeatman, J. D., Dougherty, R. F., Rykhlevskaia, E., Sherbondy, A. J., Deutsch, G. K., Wandell, B. A., & Ben-Shachar, M. (2011). Anatomical properties of the arcuate fasciculus predict phonological and reading skills in children. Journal of Cognitive Neuroscience, 23(11), 3304–3317.

Ystad, M., Hodneland, E., Adolfsdottir, S., Haász, J., Lundervold, A. J., Eichele, T., & Lundervold, A. (2011). Cortico-striatal connectivity and cognition in normal aging: a combined DTI and resting state fMRI study. NeuroImage, 55(1), 24–31.

Zelazo, P. D., Anderson, J. E., Richler, J., Wallner-Allen, K., Beaumont, J. L., & Weintraub, S. (2013). II. NIH Toolbox Cognition Battery (CB): measuring executive function and attention. Monographs of the Society for Research in Child Development, 78(4), 16–33.

