## Supplement for "Decomposing the role of alpha oscillations during brain maturation"

### Supplementary Materials

#### Section A. Rotation of the aperiodic signal

In the present analyses, a decrease of the aperiodic intercept and a flattening of the aperiodic slope were observed during brain maturation. However, these two parameters are highly correlated as a change in the aperiodic slope yields a rotation of the aperiodic signal, which in turn induces a change in the aperiodic intercept. Therefore, to investigate whether the here observed maturational decrease of the aperiodic intercept is a mere result of the flattening of the aperiodic slope, a post hoc analysis was performed. For the purpose of this analysis, the full HBN sample (N=1766) was split by the median into an older and a younger subsample. Each subsample was bootstrapped 10000 times. Pairwise, for each of the 10000 bootstrapped younger and older subsamples, the mean age difference in the aperiodic slope was calculated. Subsequently, the average aperiodic signal of each of the 10000 bootstrapped young subsamples was reconstructed by its average aperiodic intercept and slope. Each of these generated aperiodic signals was then rotated by the mean age difference to the corresponding older bootstrapped subsample. This rotation was performed using the rotate_spectrum function of the SpecParam toolbox (Donoghue et al., 2020), the rotation was centered on the mean of the power spectra (19Hz). After rotating these signals, the aperiodic intercept was extracted again using the SpecParam toolbox. This procedure yielded 10000 aperiodic intercept values, which are expected to be observed in older subjects, due to a rotation of the aperiodic signal without actual changes in the aperiodic intercept itself. These values were pairwise compared to the corresponding observed aperiodic intercept values of the bootstrapped older subjects. If no age differences would have been observed anymore, the conclusion would be that the age related change in aperiodic intercept is a mere result by the age related change in the aperiodic slope. However, after rotating the aperiodic signal of younger subjects by the age differences in aperiodic slope, the observed aperiodic intercepts were still larger than the actual aperiodic intercepts of corresponding older subjects, in all of the 10000 bootstrapped subsamples (mean age difference (older – younger) = -0.08, sd= 0.01, range= [-0.13 -0.04]). Thus, it can be concluded that there is indeed an age related decrease of the aperiodic intercept during brain maturation, which is not merely due to shifts in aperiodic slope.

#### Section B. Site-Specific Scanning Parameters

For detailled scanning protocols for all three acquisition sites see http://fcon_1000.projects.nitrc.org/indi/cmi_healthy_brain_network/ and Alexander et al. (2017).

##### *Staten Island (SI) scanning site*

Scanning at site SI was performed in a mobile trailer with a 1.5 T Siemens Avanto system and a Siemens 32-channel head coil, 32 RF receive channels and 45 mT/m gradients. The University of Minnesota Center for Magnetic Resonance Research (CMRR) simultaneous multi-slice echo planar imaging sequence was used. DTI imaging was acquired at 72 slices, with a resolution of 2 x 2 x 2 mm in 64 diffusion directions and b-values of 0, 1000 and 2000 s/mm$^2$. Further DTI parameters were TR = 3110 ms, TE = 76.2 ms, flip angle of 90 degrees and threefold multiband acceleration.

T1-weighted imaging was acquired at 176 slices with a resolution of 1 x 1 x 1 mm and scanning parameters of TR = 2730 ms, TE = 1.64 ms, TI = 1000 ms and a flip angle of 7 degrees.

##### *Rutgers University Brain Imaging Center (RUBIC) scanning site*

At the RUBIC site, scans were acquired with a Siemens 3T Tim Trio MRI scanner, a Siemens 32-channel head coil using the CMRR simultaneous multi-slice echo planar imaging sequence. DTI imaging comprised 72 slices with a resolution of 1.8 x 1.8 x 1.8 mm. Further DTI specifications were: TR = 3320 ms, TE = 100.2 ms, a flip angle of 90 degrees, threefold multiband acceleration, 64 diffusion directions and b-values of 0, 1000 and 2000 s/mm$^2$.

For T1-weighted imaging 224 slices at a resolution of 0.8 x 0.8 x 0.8 mm were acquired with TR = 2500 ms, TE = 3.15 ms, TI = 1060 ms and flip angle of 8 degrees.

##### *CitiGroup Cornell Brain Imaging Center (CBIC) scanning sites*

The CBIC site was equipped with a 3T Prisma scanner and a Siemens 32-channel head coil using the CMRR simultaneous multi-slice echo planar imaging sequence. DTI was acquired at 81 slices with a resolution of 1.8 x 1.8 x 1.8 mm in 64 diffusion directions and b-values of 0, 1000 and 2000 s/mm$^2$. Further scanning parameters were set to TR = 3320 ms, TE = 100.2 ms, a flip angle of 90 deg, threefold multiband acceleration. The parameters for T1-weighted imaging at CBIC were identical to the RUBIC site.

#### Section C. DTI Preprocessing

The DTI preprocessing pipeline included the following steps: (1) Denoising was performed using the “MPdenoising” function employing a 4-dimensional image denoising and noise map estimation algorithm (Veraart, Fieremans, & Novikov, 2016). (2) Gibbs artefacts were removed from the denoised images using the “unring” function with the default parameters (Kellner, Dhital, Kiselev, & Reisert, 2016). The following steps were executed using the FMRIB Software Library (FSL) version 6.0.4 (Jenkinson, Beckmann, Behrens, Woolrich, & Smith, 2012). (3) We used the FSL Brain Extraction Tool (BET) to obtain a binary brain mask from the whole head image using a fractional anisotropy threshold of 0.1 (Smith, 2002). (4) Eddy current-induced distortions and in-scanner head motion artefacts were removed using the FSL tool “eddy_cuda” (Andersson et al., 2017; Andersson, Graham, Zsoldos, & Sotiropoulos, 2016; Andersson & Sotiropoulos, 2016). For this tool the settings were as follows: 8 iterations, smoothing full-width-half-max parameters of [10, 6, 4, 2, 0, 0, 0, 0] for each respective iteration, enabled outlier detection and replacement for slice-wise and multiband group outliers and additional 8 iterations for slice-to-volume correction. (5) Outlier detection of MRI parameters and robust parameter estimation was performed with the function “irlls” applying iterative reweighted linear least squares (Collier, Veraart, Jeurissen, Dekker, & Sijbers, 2015). (6) Using the function “dki_fit” (Veraart, Sijbers, Sunaert, Leemans, & Jeurissen, 2013) we performed tensor fitting and extraction of diffusivity measures based on weighted linear least squares estimation. (7) For the extraction of DTI parameters we used the function “dki_parameters” (Veraart et al., 2011). (8) To calculate white matter tract integrity metrics we applied the function “wmti_parameters” (Fieremans, Jensen, & Helpern, 2011). (8) AC-PC aligned nifti image was created using the function “mrAnatAutoAlignAcpcNifti” (https://github.com/vistalab/vistasoft, Vistalab, Stanford University, Stanford, CA). The output of this preprocessing pipeline was the input for tractography using Automating Fiber-Tract Quantification (AFQ, v0.1) (Yeatman, Dougherty, Myall, Wandell, & Feldman, 2012). (9) The parameters for the function “AFQ_Create” were: “cutoff” of [5 95], FA threshold of 0.2, FA mask threshold of 0.3, and angle threshold of 35 degrees. (10) We used the “AFQ_run” function to calculate individual white matter tract profiles of FA. (11) Based on these tract profiles we calculated a mean FA value which is the average FA of 100 equidistant nodes along clipped predefined regions of the left and the right thalamic radiation respectively.

#### Section D. T1-weighted Preprocessing

T1-weighted scans were preprocessed with FreeSurfer (version 6.0.0) (http://surfer.nmr.mgh.harvard.edu/). Subcortical volumetric segmentation of the left and right thalamus was computed using the function recon-all (Fischl et al., 2002; Fischl et al., 2004; Han et al., 2006). First, the T1-weighted scan was corrected for intensity non-uniformity with Non-parametric Non-uniform intensity Normalization. Second, the Talairach transformation was computed which is an affine transform to the MNI305 atlas. Third, intensity normalization was applied, which corrects deviations in intensity for later intensity-based segmentation. All voxel's intensity was scaled such that the white matter mean intensity was 110. Fourth, skull tissue was removed from the intensity normalized image and a brain mask was created. Next automatic subcortical segmentation was performed. For this, the intensity non-uniformity corrected (NU-corrected) image was co-registered to the Gaussian Classifier Atlas (GCA). Based on the GCA model the image was first normalized and nonlinearly transformed to the GCA atlas. In a next step, regions corresponding to the neck were removed from the NU-corrected image. Further, a transformation was applied of the NU-corrected image without the neck to the GCA image containing the skull. Lastly, subcortical structures were labeled in accordance with the GCA model and statistics on the segmented subcortical structures computed and summarized.

#### Section E. Simulating confounds in relative alpha power

To highlight confounding factors in the analysis of age related changes in relative alpha power, simulations were performed. Therefore, power spectra were simulated as defined in the specparam algorithm (see equation 1 and 2, section 2.2.6). Gaussians representing neural oscillations were further defined as in Donoghue et al. (2020):

$$G_n = a \times \exp\left(\frac{-(F-c)^2}{2w^2}\right)$$

First, two power spectra were generated sharing the identical aperiodic signal component (intercept =1.82, exponent =1.99). The same alpha oscillation was added to both power spectra, being defined by the center frequency c=10.5 Hz, the power a = $\log_{10}(6)$, and the bandwidth w=1. A larger theta oscillation as added to the first power spectrum (simulated data 1, c=4, a= $\log_{10}(7.5)$ and w=1.1) than to the second power spectrum (simulated data 2, c=4, a= $\log_{10}(2)$ and w=1.1). Resulting power spectra are plotted in supplementary Figure 4A.

For the second simulated case, two power spectra were generated, which differed in the aperiodic signal component (simulated data 3: intercept=1.8, exponent =2.0; simulated data 4: intercept =1.5, exponent= 1.8). The same alpha oscillation was added to both simulated power spectra (c=10, a= $\log_{10}(6)$ and w=1). Resulting power spectra are visualized in supplementary Figure 4B. For both scenarios, differences in relative and total individualized alpha power were calculated.

#### Section F. Relation of alpha power to Flanker task scores

To investigate the relationship between the different measures of alpha power and attentional performance, the flanker task of the National Institutes of Health Toolbox Cognition Battery was employed. The total Flanker score was used as the predictor in the linear models. The univariate linear models controlled for age, gender and handedness and were defined as:

$$alpha\ power \sim Flanker\ score +\ age + gender + EHQ$$

The models were fitted once for the outcome variable total individualized alpha power, and once for aperiodic-adjusted individualized alpha power.

#### Supplementary Figure 1

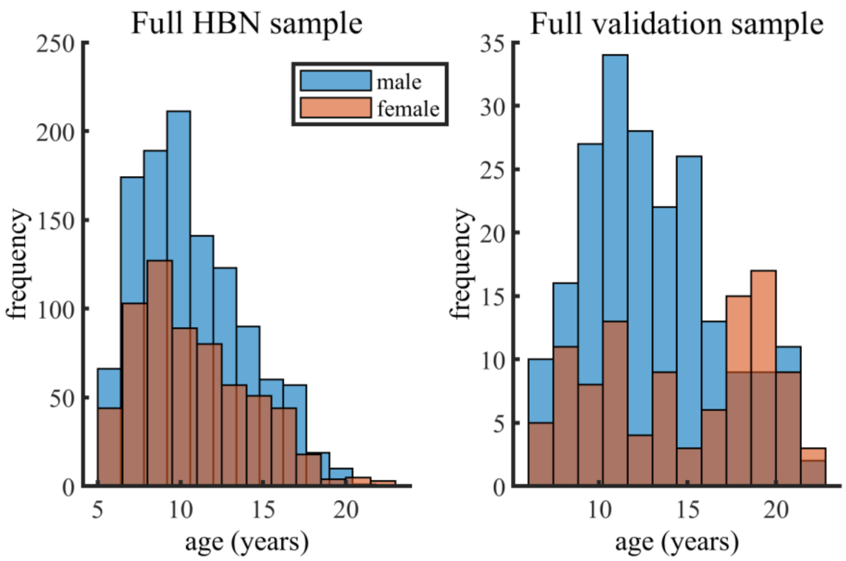


Distribution of age and gender in the final included samples

#### Supplementary Figure 2

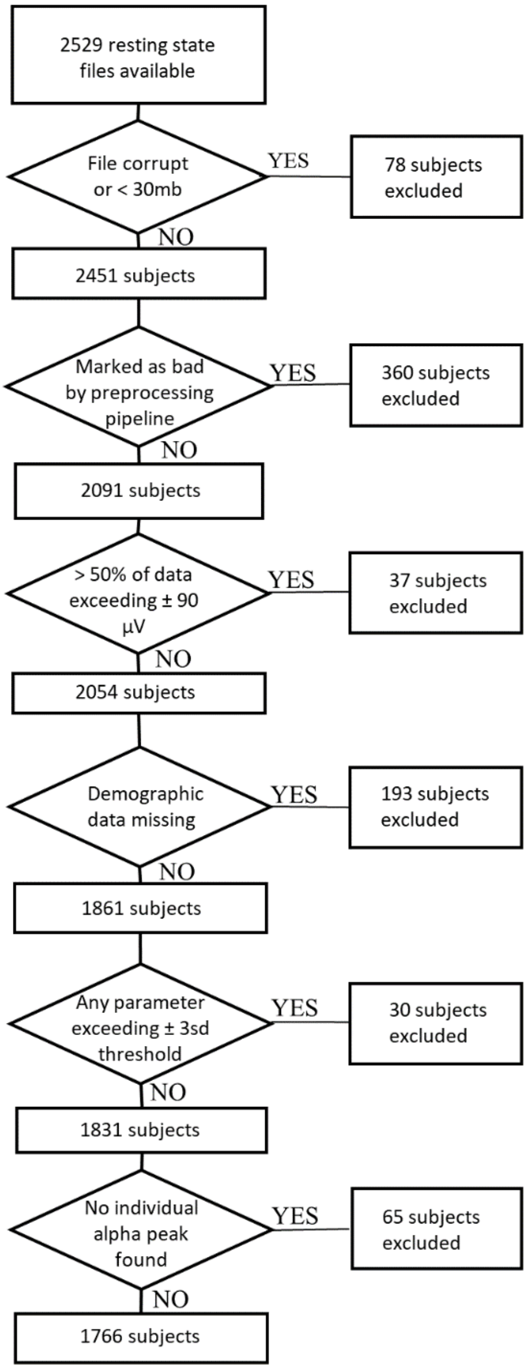


Flow chart of exclusion criteria applied to the main HBN dataset.

#### Supplementary Figure 3

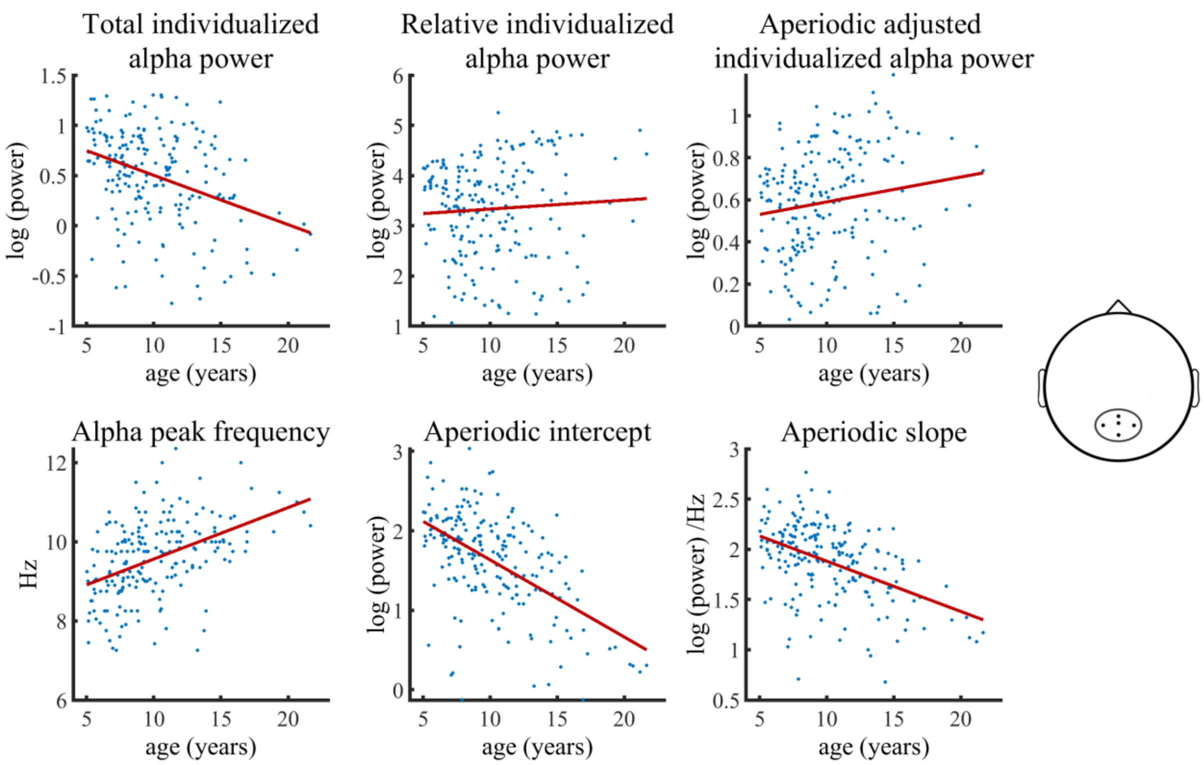


Visualization of data of the HBN subsample without any given diagnosis, as used in the Bayesian regression models. Solid lines represent fitted regression lines. The schematic head (right) indicates the location of the electrode cluster, of which data was aggregated.

#### Supplementary Figure 4

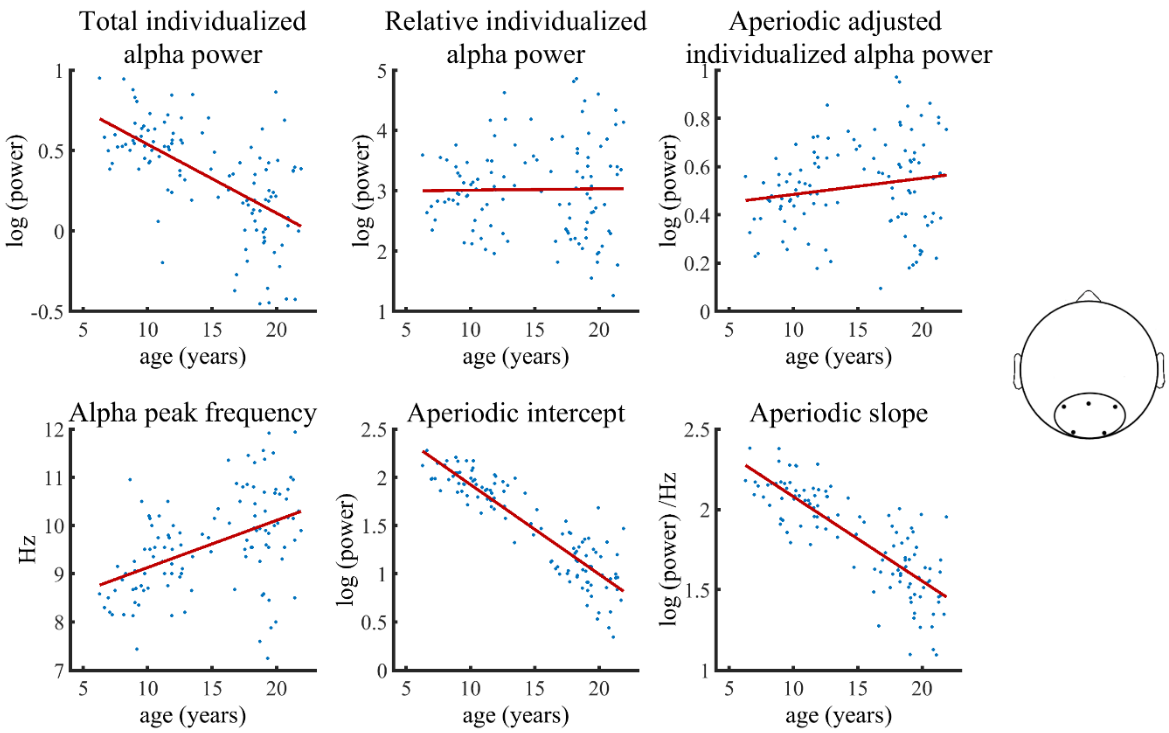


Visualization of data of the healthy validation subsample used in the Bayesian regression model. Solid lines represent fitted regression lines. The schematic head on the top right indicates the location of the electrode cluster, of which data was aggregated.

#### Supplementary Figure 5

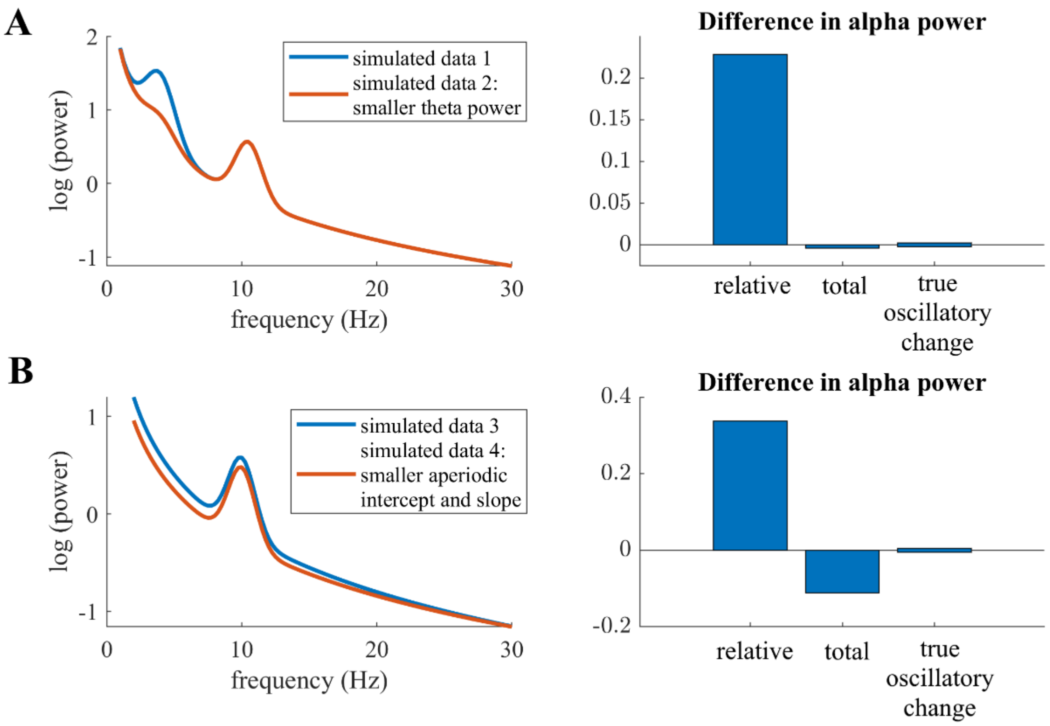


Visualizations of possible fallacies in relative power measures in simulated data. Bar plots on the right indicate the difference in alpha power between simulated data 2 and simulated data 1 in A and simulated data 4 to simulated data 3 in B. A: Two simulated power spectra with identical true alpha oscillatory power. The high amplitude oscillation in the theta range (~5Hz) in simulated data 1 conflates results in relative power differences in the alpha band. B: Two simulated power spectra with identical alpha oscillatory power. Here, differences in aperiodic intercept and slope between the two signals conflate results in relative power differences in the alpha band.

#### Supplementary Table 1

*Model comparison*

| | WAIC | ELDP_diff (se) |
|---|---|---|
| Subsample without any given diagnosis | | |
| $[dv's] \sim age + gender + EHQ + site$ | -183.03 | 0.00 (0.00) |
| $[dv's] \sim age * gender + EHQ + site$ | -172.19 | -5.42 (4.02) |
| Full Sample, categorical diagnosis variable | | |
| $[dv's] \sim age + gender + diagnosis + EHQ + site$ | -1534.40 | -19.70 (7.98) |
| $[dv's] \sim age * gender + diagnosis + EHQ + site$ | -1573.79 | 0.00 (0.00) |
| $[dv's] \sim age + gender * diagnosis + EHQ + site$ | -1526.36 | -23.72 (8.16) |
| $[dv's] \sim age * diagnosis + age * gender + EHQ + site$ | -1561.44 | -6.18 (2.99) |
| $[dv's] \sim age * gender * diagnosis + EHQ + site$ | -1559.82 | -6.99 (8.42) |

*Note:* Dependent variables (dv's) used for the model comparisons are : Total individualized alpha power, aperiodic-adjusted individualized alpha power, relative individualized alpha power, aperiodic intercept, aperiodic slope and IAF. WAIC refers to the Watanabe Akaike (or: widely applicable) information criterion. The difference of the expected log pointwise predictive density (ELDP, i.e. the expected predictive accuracy of the model) to the best model is calculated additionally, together with the standard error of this difference.

#### Supplementary Table 2

*Bayesian regression model results of HBN subsample of subjects without any given diagnosis*

| | $\beta_{predictor}$ [CI] | |
|---|---|---|
| Outcome | age | gender |
| Alpha peak frequency | 0.40 [0.22 .58] | -0.11 [-0.31 0.08] |
| Total individualized alpha power | -0.21 [-0.40 -0.03] | -0.38 [-0.57 -0.21] |
| Relative individualized alpha power | 0.10 [-0.10 0.30] | -0.25 [-0.46 -0.06] |
| Aperiodic-adjusted individualized alpha power | 0.21 [0.02 0.41] | -0.32 [-0.53 -0.13] |
| Aperiodic intercept | -0.42 [-0.59 -0.25] | -0.37 [-0.55 -0.21] |
| Aperiodic slope | -0.38 [-0.55 -0.20] | -0.33 [-0.51 -0.17] |

*Note:* CI = 99.17% Credible Interval, gender variable is coded as: 1=female, 0=male.

#### Supplementary Table 3

*Bayesian regression model results of full HBN sample with categorical sub-diagnosis predictor*

| | | | $\beta_{predictor}$ [CI] | | | |
|---|---|---|---|---|---|---|
| Outcome | age | gender | diagnosis: ADHD (inattentive) | diagnosis: ADHD (combined) | diagnosis: Other | age*gender |
| alpha peak frequency | 0.41 [0.33 0.49] | -0.09 [-0.15 -0.03] | -0.07 [-0.18 0.03] | -0.08 [-0.19 0.03] | -0.07 [-0.18 0.04] | -0.02 [-0.14 0.10] |
| total individualized alpha power | -0.31 [-0.39 -0.24] | -0.36 [-0.43 -0.30] | 0.02 [-0.09 0.12] | 0.01 [-0.09 0.12] | 0.01 [-0.09 0.11] | 0.13 [0.01 0.25] |
| Relative individualized alpha power | 0.14 [0.06 0.23] | -0.33 [-0.40 -0.27] | -0.00 [-0.11 0.11] | -0.01 [-0.12 0.10] | 0.01 [-0.10 0.12] | -0.05 [-0.18 0.08] |
| aperiodic-adjusted individualized alpha power | 0.23 [0.15 0.30] | -0.38 [-0.44 -0.32] | -0.02 [-0.13 0.08] | -0.05 [-0.16 0.05] | -0.03 [-0.13 0.08] | -0.05 [-0.17 0.08] |
| aperiodic intercept | -0.54 [-0.62 -0.48] | -0.36 [-0.42 -0.31] | -0.01 [-0.11 0.08] | 0.00 [-0.10 0.09] | -0.02 [-0.11 0.08] | 0.08 [-0.03 0.19] |
| aperiodic slope | -0.45 [-0.52 -0.38] | -0.38 [-0.43 -0.32] | -0.04 [-0.13 0.06] | -0.03 [-0.13 0.07] | -0.03 [-0.13 0.06] | -0.05 [-0.16 0.06] |

*Note:* CI = 99.17% Credible Interval

#### Supplementary Table 4

*Bayesian regression model results for canonical alpha power measures in the full HBN sample.*

| | $\beta_{predictor}$ [CI] | | | | |
|---|---|---|---|---|---|
| Outcome | age | gender | diagnosis: ADHD | diagnosis: Other | age*gender |
| total canonical alpha power | -0.10 [-0.17 - 0.02] | -0.39 [-0.45 - 0.33] | -0.02 [-0.12 0.08] | -0.01 [-0.12 0.10] | 0.11 [-0.01 0.23] |
| Relative canonical alpha power | 0.36 [0.29 0.43] | -0.32 [-0.38 - 0.26] | -0.03 [-0.13 0.06] | -0.01 [-0.11 0.09] | -0.07 [-0.18 0.05] |
| aperiodic-adjusted canonical alpha power | 0.31 [0.24 0.39] | -0.38 [-0.44 - 0.32] | -0.05 [-0.15 0.04] | -0.02 [-0.13 0.08] | -0.07 [-0.18 0.04] |

*Note:* CI = 99.17% Credible Interval. Due to model convergence problems, models were multivariately estimated using either total, relative or aperiodic-adjusted canonical alpha power as dependent variable, each together with alpha peak frequency, aperiodic intercept and the aperiodic slope. Alpha peak frequency, aperiodic intercept and periodic slope were only added as outcome measures to account for correlations among periodic and aperiodic measures, but resulting parameter estimates did not change compared to analysis done on the same dataset using individualized alpha band measures (see 3.1, Table 3). Therefore, they are omitted here.

#### Supplementary Table 5

*Validation study: Bayesian regression model results of subjects without any given diagnosis, using uninformative priors.*

| | $\beta_{predictor}$ [CI] | |
|---|---|---|
| Outcome | age | gender |
| Alpha peak frequency | 0.44 [0.21 .68] | -0.12 [-0.36 0.12] |
| Total individualized alpha power | -0.60 [-0.80 -0.40] | -0.02 [-0.24 -0.18] |
| Relative individualized alpha power | 0.08 [-0.19 0.34] | 0.02 [-0.25 0.29] |
| Aperiodic-adjusted individualized alpha power | 0.24 [-0.03 0.50] | -0.05 [-0.31 0.21] |
| Aperiodic intercept | -0.88 [-1.00 -0.76] | -0.06 [-0.17 0.05] |
| Aperiodic slope | -0.80 [-0.96 -0.64] | -0.05 [-0.21 0.10] |

*Note:* CI = 98.97% Credible Interval, gender variable is dummy coded: 1=female, 0=male.

#### Supplementary Table 6

*Validation study: Bayesian regression model results using informative priors extracted from the main HBN analysis.*

| | | | $\beta_{predictor}$ [CI] | |
|---|---|---|---|---|
| Outcome | age | gender | diagnosis: ADHD | age*gender |
| Alpha peak frequency | 0.38 [0.32 .44] | -0.06 [-0.11 0.00] | -0.08 [-0.16 0.00] | -0.05 [-0.15 0.05] |
| Total individualized alpha power | -0.35 [-0.41 -0.30] | -0.38 [-0.43 -0.34] | -0.01 [-0.05 0.08] | 0.06 [-0.02 0.15] |
| Relative individualized alpha power | 0.13 [0.08 0.19] | -0.29 [-0.34 -0.24] | -0.03 [-0.10 0.04] | -0.02 [-0.11 0.07] |
| Aperiodic-adjusted individualized alpha power | 0.31 [0.25 0.36] | -0.39 [-44 -0.35] | -0.07 [-0.14 -0.01] | 0.00 [-0.08 0.08] |
| Aperiodic intercept | -0.60 [-0.64 -0.56] | -0.36 [-0.39 -0.32] | -0.01 [-0.06 0.03] | 0.01 [-0.05 0.08] |
| Aperiodic slope | -0.46 [-0.51 -0.41] | -0.37 [-0.42 -0.33] | -0.06 [-0.12 0.00] | -0.06 [-0.14 0.03] |

*Note:* CI = 98.97% Credible Interval, gender variable is dummy coded: 1=female, 0=male.

#### Supplementary table 7

*Bayesian regression models investigating influences f white matter integrity of the left and right thalamic radiations as well as global white matter on aperiodic signal parameters in the HBN dataset.*

| | $\beta_{predictor}$ [CI] | | |
|---|---|---|---|
| Outcome | left thalamic radiation | right thalamic radiation | global white matter |
| Aperiodic intercept | 0.04 [-0.03 0.11] | 0.01 [-0.06 0.08] | 0.02 [-0.05 0.10] |
| Aperiodic slope | 0.04 [-0.01 0.09] | 0.02 [-0.03 0.07] | 0.03 [-0.02 0.08] |

*Note:* CI = 95% Credible Interval. Due to collinearity between white matter integrity of the left and right thalamic radiations, multivariate models were fitted for either the predictor left thalamic radiation, right thalamic radiation or global white matter integrity on the aperiodic signal intercept and slope, while controlling for age, gender and total intracranial volume.

#### Supplementary table 8

*Linear models determining the influence of Flanker total task scores on total and aperiodic-adjusted individualized alpha power.*

| | $\beta_{predictor}$ (standard error) | | |
|---|---|---|---|
| Outcome | Flanker total score | age | gender |
| Aperiodic-adjusted individualized alpha power | 0.001 (0.0003) ** | 0.01 (0.001) *** | -0.18 (0.011) *** |
| Total individualized alpha power | 0.001 (0.0006) $^{n.s.}$ | -0.03 (0.003) *** | -0.32 (0.019)*** |

*Note:* ** indicates p values < 0.01, *** indicates p values < 0.001, n.s.= not significant

### References


Ades-Aron, B., Veraart, J., Kochunov, P., McGuire, S., Sherman, P., Kellner, E., . . . Fieremans, E. (2018). Evaluation of the accuracy and precision of the diffusion parameter EStImation with Gibbs and NoisE removal pipeline. *NeuroImage*, *183*, 532–543. https://doi.org/10.1016/j.neuroimage.2018.07.066

Alcauter, S., Lin, W., Smith, J. K., Short, S. J., Goldman, B. D., Reznick, J. S., . . . Gao, W. (2014). Development of thalamocortical connectivity during infancy and its cognitive correlations. *Journal of Neuroscience*, *34*(27), 9067–9075. https://doi.org/10.1523/JNEUROSCI.0796-14.2014

Alexander, L. M., Escalera, J., Ai, L., Andreotti, C., Febre, K., Mangone, A., . . . Milham, M. P. (2017). An open resource for transdiagnostic research in pediatric mental health and learning disorders. *Scientific Data*, *4*, 170181. https://doi.org/10.1038/sdata.2017.181

Alkonyi, B., Juhász, C., Muzik, O., Behen, M. E., Jeong, J.-W., & Chugani, H. T. (2011). Thalamocortical connectivity in healthy children: Asymmetries and robust developmental changes between ages 8 and 17 years. *American Journal of Neuroradiology*, *32*(5), 962–969. https://doi.org/10.3174/ajnr.A2417

Andersson, J. L. R., Graham, M. S., Drobnjak, I., Zhang, H., Filippini, N., & Bastiani, M. (2017). Towards a comprehensive framework for movement and distortion correction of diffusion MR images: Within volume movement. *NeuroImage*, *152*, 450–466. https://doi.org/10.1016/j.neuroimage.2017.02.085

Andersson, J. L. R., Graham, M. S., Zsoldos, E., & Sotiropoulos, S. N. (2016). Incorporating outlier detection and replacement into a non-parametric framework for movement and distortion correction of diffusion MR images. *NeuroImage*, *141*, 556–572. https://doi.org/10.1016/j.neuroimage.2016.06.058

Andersson, J. L. R., & Sotiropoulos, S. N. (2016). An integrated approach to correction for off-resonance effects and subject movement in diffusion MR imaging. *NeuroImage*, *125*, 1063–1078. https://doi.org/10.1016/j.neuroimage.2015.10.019

Babiloni, C., Barry, R. J., Başar, E., Blinowska, K. J., Cichocki, A., Drinkenburg, W. H. I. M., . . . Hallett, M. (2020). International Federation of Clinical Neurophysiology (IFCN) - EEG research workgroup: Recommendations on frequency and topographic analysis of resting state EEG rhythms. Part 1: Applications in clinical research studies. *Clinical Neurophysiology : Official Journal of the International Federation of Clinical Neurophysiology*, *131*(1), 285–307. https://doi.org/10.1016/j.clinph.2019.06.234

Ball, G., Pazderova, L., Chew, A., Tusor, N., Merchant, N., Arichi, T., . . . Counsell, S. J. (2015). Thalamocortical Connectivity Predicts Cognition in Children Born Preterm. *Cerebral Cortex (New York, N.Y. : 1991)*, *25*(11), 4310–4318. https://doi.org/10.1093/cercor/bhu331

Basser, P. J., Pajevic, S., Pierpaoli, C., Duda, J., & Aldroubi, A. (2000). In vivo fiber tractography using DT-MRI data. *Magnetic Resonance in Medicine*, *44*(4), 625–632. https://doi.org/10.1002/1522-2594(200010)44:4<625::AID-MRM17>3.0.CO;2-O

Bazanova, O. M., & Vernon, D. (2014). Interpreting EEG alpha activity. *Neuroscience and Biobehavioral Reviews*, *44*, 94–110. https://doi.org/10.1016/j.neubiorev.2013.05.007

Benninger, C. [C.], Matthis, P., & Scheffner, D. (1984). EEG development of healthy boys and girls. Results of a longitudinal study. *Electroencephalography and Clinical Neurophysiology*, *57*(1), 1–12. https://doi.org/10.1016/0013-4694(84)90002-6

Bishop, G. H. (1936). The INTERPRETATION OF CORTICAL POTENTIALS. *Cold Spring Harbor Symposia on Quantitative Biology*, *4*(0), 305–319. https://doi.org/10.1101/SQB.1936.004.01.032

Bürkner, P.-C. (2017, May 31). *Advanced Bayesian Multilevel Modeling with the R Package brms*. Retrieved from http://arxiv.org/pdf/1705.11123v2

Carlson, H. L., Laliberté, C., Brooks, B. L., Hodge, J., Kirton, A., Bello-Espinosa, L., . . . Sherman, E. M. S. (2014). Reliability and variability of diffusion tensor imaging (DTI) tractography in pediatric epilepsy. *Epilepsy & Behavior : E&B*, *37*, 116–122. https://doi.org/10.1016/j.yebeh.2014.06.020

Cellier, D., Riddle, J., Petersen, I., & Hwang, K. (2021). The development of theta and alpha neural oscillations from ages 3 to 24 years. *Developmental Cognitive Neuroscience*, *50*, 100969. https://doi.org/10.1016/j.dcn.2021.100969

Charlton, R. A., Barrick, T. R., Lawes, I. N. C., Markus, H. S., & Morris, R. G. (2010). White matter pathways associated with working memory in normal aging. *Cortex; a Journal Devoted to the Study of the Nervous System and Behavior*, *46*(4), 474–489. https://doi.org/10.1016/j.cortex.2009.07.005

Cheveigné, A. de (2020). Zapline: A simple and effective method to remove power line artifacts. *NeuroImage*, *207*, 116356. https://doi.org/10.1016/j.neuroimage.2019.116356

Clarke, A. R., Barry, R. J., McCarthy, R., & Selikowitz, M. (2001). Age and sex effects in the EEG: development of the normal child. *Clinical Neurophysiology*, *112*(5), 806–814. https://doi.org/10.1016/S1388-2457(01)00488-6

Collier, Q., Veraart, J., Jeurissen, B., Dekker, A. J. den, & Sijbers, J. (2015). Iterative reweighted linear least squares for accurate, fast, and robust estimation of diffusion magnetic resonance parameters. *Magnetic Resonance in Medicine*, *73*(6), 2174–2184. https://doi.org/10.1002/mrm.25351

Cragg, L., Kovacevic, N., McIntosh, A. R. [Anthony Randal], Poulsen, C., Martinu, K., Leonard, G., & Paus, T. [Tomáš] (2011). Maturation of EEG power spectra in early adolescence: A longitudinal study. *Developmental Science*, *14*(5), 935–943. https://doi.org/10.1111/j.1467-7687.2010.01031.x

Delignette-Muller, M. L., & Dutang, C. (2015). Fitdistrplus : An R Package for Fitting Distributions. *Journal of Statistical Software*, *64*(4). https://doi.org/10.18637/jss.v064.i04

Delorme, A., & Makeig, S. (2004). Eeglab: An open source toolbox for analysis of single-trial EEG dynamics including independent component analysis. *Journal of Neuroscience Methods*, *134*(1), 9–21. https://doi.org/10.1016/j.jneumeth.2003.10.009

Díaz de León, A. E., Harmony, T. [T.], Marosi, E. [E.], Becker, J. [J.], & Alvarez, A. (1988). Effect of different factors on EEG spectral parameters. *The International Journal of Neuroscience*, *43*(1-2), 123–131. https://doi.org/10.3109/00207458808985789

Donoghue, T., Dominguez, J., & Voytek, B. [Bradley] (2020). Electrophysiological Frequency Band Ratio Measures Conflate Periodic and Aperiodic Neural Activity. *ENeuro*. Advance online publication. https://doi.org/10.1523/ENEURO.0192-20.2020

Donoghue, T., Haller, M., Peterson, E., Varma, P., Sebastian, P., Gao, R. [R.], . . . Voytek, B. [B.] (2020). Parameterizing neural power spectra into periodic and aperiodic components. *Nature Neuroscience*, *in press*.

Dustman, R. E., Shearer, D. E., & Emmerson, R. Y. (1999). Life-span changes in EEG spectral amplitude, amplitude variability and mean frequency. *Clinical Neurophysiology*, *110*(8), 1399–1409. https://doi.org/10.1016/S1388-2457(99)00102-9

Fair, D. A., Bathula, D., Mills, K. L., Dias, T. G. C., Blythe, M. S., Zhang, D., . . . Nagel, B. J. (2010). Maturing thalamocortical functional connectivity across development. *Frontiers in Systems Neuroscience*, *4*, 10. https://doi.org/10.3389/fnsys.2010.00010

Fieremans, E., Jensen, J. H., & Helpern, J. A. (2011). White matter characterization with diffusional kurtosis imaging. *NeuroImage*, *58*(1), 177–188. https://doi.org/10.1016/j.neuroimage.2011.06.006

Fischl, B., Salat, D. H., Busa, E., Albert, M., Dieterich, M., Haselgrove, C., . . . Dale, A. M. (2002). Whole Brain Segmentation. *Neuron*, *33*(3), 341–355. https://doi.org/10.1016/S0896-6273(02)00569-X

Fischl, B., Salat, D. H., van der Kouwe, A. J. W., Makris, N., Ségonne, F., Quinn, B. T., & Dale, A. M. (2004). Sequence-independent segmentation of magnetic resonance images. *NeuroImage*, *23 Suppl 1*, S69-84. https://doi.org/10.1016/j.neuroimage.2004.07.016

Forman, A. K., Poswanger, K., & Waldherr, K. (2006). *Grundintelligenztest Skala 2 (CFT 20-R) mit Wortschatztest (WS) und Zahlenfolgentest (ZF).*

Foxe, J. J., & Snyder, A. C. (2011). The Role of Alpha-Band Brain Oscillations as a Sensory Suppression Mechanism during Selective Attention. *Frontiers in Psychology*, *2*, 154. https://doi.org/10.3389/fpsyg.2011.00154

Gao, R. [Richard], Peterson, E. J., & Voytek, B. [Bradley] (2017). Inferring synaptic excitation/inhibition balance from field potentials. *NeuroImage*, *158*, 70–78. https://doi.org/10.1016/j.neuroimage.2017.06.078

Gasser, T., Verleger, R., Bächer, P., & Sroka, L. (1988). Development of the EEG of school-age children and adolescents. I. Analysis of band power. *Electroencephalography and Clinical Neurophysiology*, *69*(2), 91–99. https://doi.org/10.1016/0013-4694(88)90204-0

Gelman, A., Jakulin, A., Su, Y.-S., & Pittau, M. G. (2007). A Default Prior Distribution for Logistic and Other Regression Models. *SSRN Electronic Journal.* Advance online publication. https://doi.org/10.2139/ssrn.1010421

Giedd, J. N., Blumenthal, J., Jeffries, N. O., Castellanos, F. X. [F. X.], Liu, H., Zijdenbos, A., . . . Rapoport, J. L. (1999). Brain development during childhood and adolescence: A longitudinal MRI study. *Nature Neuroscience*, *2*(10), 861–863. https://doi.org/10.1038/13158

Gómez, C. M., Rodríguez-Martínez, E. I., Fernández, A., Maestú, F., Poza, J., & Gómez, C. (2017). Absolute Power Spectral Density Changes in the Magnetoencephalographic Activity During the Transition from Childhood to Adulthood. *Brain Topography*, *30*(1), 87–97. https://doi.org/10.1007/s10548-016-0532-0

Han, X., Jovicich, J., Salat, D., van der Kouwe, A., Quinn, B., Czanner, S., . . . Fischl, B. (2006). Reliability of MRI-derived measurements of human cerebral cortical thickness: The effects of field strength, scanner upgrade and manufacturer. *NeuroImage*, *32*(1), 180–194. https://doi.org/10.1016/j.neuroimage.2006.02.051

Harmony, T. [Thalía], Marosi, E. [Erzsébet], Becker, J. [Jacqueline], Rodríguez, M., Reyes, A., Fernández, T., . . . Bernal, J. (1995). Longitudinal quantitative EEG study of children with different performances on a reading-writing test. *Electroencephalography and Clinical Neurophysiology*, *95*(6), 426–433. https://doi.org/10.1016/0013-4694(95)00135-2

He, W., Donoghue, T., Sowman, P. F., Seymour, R. A., Brock, J., Crain, S., . . . Hillebrand, A. (2019). *Co-Increasing Neuronal Noise and Beta Power in the Developing Brain.* https://doi.org/10.1101/839258

Hua, K., Zhang, J., Wakana, S., Jiang, H., Li, X., Reich, D. S., . . . Mori, S. (2008). Tract probability maps in stereotaxic spaces: Analyses of white matter anatomy and tract-specific quantification. *NeuroImage*, *39*(1), 336–347. https://doi.org/10.1016/j.neuroimage.2007.07.053

Hughes, A. M., Whitten, T. A., Caplan, J. B., & Dickson, C. T. (2012). Bosc: A better oscillation detection method, extracts both sustained and transient rhythms from rat hippocampal recordings. *Hippocampus*, *22*(6), 1417–1428. https://doi.org/10.1002/hipo.20979

Hughes, E. J., Bond, J., Svrckova, P., Makropoulos, A., Ball, G., Sharp, D. J., . . . Counsell, S. J. (2012). Regional changes in thalamic shape and volume with increasing age. *NeuroImage*, *63*(3), 1134–1142. https://doi.org/10.1016/j.neuroimage.2012.07.043

Huttenlocher, P. R., & Courten, C. de (1987). The development of synapses in striate cortex of man. *Human Neurobiology*, *6*(1), 1–9.

Jenkinson, M., Beckmann, C. F., Behrens, T. E. J., Woolrich, M. W., & Smith, S. M. (2012). Fsl. *NeuroImage*, *62*(2), 782–790. https://doi.org/10.1016/j.neuroimage.2011.09.015

Jin, Y., O'Halloran, J. P., Plon, L., Sandman, C. A., & Potkin, S. G. (2006). Alpha EEG predicts visual reaction time. *The International Journal of Neuroscience*, *116*(9), 1035–1044. https://doi.org/10.1080/00207450600553232

John, E. R., Ahn, H., Prichep, L., Trepetin, M., Brown, D., & Kaye, H. (1980). Developmental equations for the electroencephalogram. *Science (New York, N.Y.)*, *210*(4475), 1255–1258. https://doi.org/10.1126/science.7434026

Kail, R. (2000). Speed of Information Processing. *Journal of School Psychology*, *38*(1), 51–61. https://doi.org/10.1016/S0022-4405(99)00036-9

Kaufman, J., Birmaher, B., Brent, D., Rao, U., Flynn, C., Moreci, P., . . . Ryan, N. (1997). Schedule for Affective Disorders and Schizophrenia for School-Age Children-Present and Lifetime Version (K-SADS-PL): Initial reliability and validity data. *Journal of the American Academy of Child & Adolescent Psychiatry*, *36*(7), 980–988. https://doi.org/10.1097/00004583-199707000-00021

Keller, S. S., Gerdes, J. S., Mohammadi, S., Kellinghaus, C., Kugel, H., Deppe, K., . . . Deppe, M. (2012). Volume estimation of the thalamus using freesurfer and stereology: Consistency between methods. *Neuroinformatics*, *10*(4), 341–350. https://doi.org/10.1007/s12021-012-9147-0

Kellner, E., Dhital, B., Kiselev, V. G., & Reisert, M. (2016). Gibbs-ringing artifact removal based on local subvoxel-shifts. *Magnetic Resonance in Medicine*, *76*(5), 1574–1581. https://doi.org/10.1002/mrm.26054

Kessler, R. C., Berglund, P., Demler, O., Jin, R., Merikangas, K. R., & Walters, E. E. (2005). Lifetime prevalence and age-of-onset distributions of DSM-IV disorders in the National Comorbidity Survey Replication. *Archives of General Psychiatry*, *62*(6), 593–602. https://doi.org/10.1001/archpsyc.62.6.593

Klimesch, W. [W.], Doppelmayr, M., Schimke, H., & Pachinger, T. (1996). Alpha Frequency, Reaction Time, and the Speed of Processing Information. *Journal of Clinical Neurophysiology*, *13*(6), 511. Retrieved from https://journals.lww.com/clinicalneurophys/fulltext/1996/11000/alpha_frequency,_reaction_time,_and_the_speed_of.6.aspx

Klimesch, W. [Wolfgang] (1997). EEG-alpha rhythms and memory processes. *International Journal of Psychophysiology*, *26*(1-3), 319–340. https://doi.org/10.1016/S0167-8760(97)00773-3

Klimesch, W. [Wolfgang] (1999). EEG alpha and theta oscillations reflect cognitive and memory performance: a review and analysis. *Brain Research Reviews*, *29*(2-3), 169–195. https://doi.org/10.1016/S0165-0173(98)00056-3

Klimesch, W. [Wolfgang] (2012). A-band oscillations, attention, and controlled access to stored information. *Trends in Cognitive Sciences*, *16*(12), 606–617. https://doi.org/10.1016/j.tics.2012.10.007

Langer, N., Bastian, C. C. von, Wirz, H., Oberauer, K., & Jäncke, L. (2013). The effects of working memory training on functional brain network efficiency. *Cortex; a Journal Devoted to the Study of the Nervous System and Behavior*, *49*(9), 2424–2438. https://doi.org/10.1016/j.cortex.2013.01.008

Langer, N., Pedroni, A., Gianotti, L. R. R., Hänggi, J., Knoch, D., & Jäncke, L. (2012). Functional brain network efficiency predicts intelligence. *Human Brain Mapping*, *33*(6), 1393–1406. https://doi.org/10.1002/hbm.21297

Laufs, H., Holt, J. L., Elfont, R., Krams, M., Paul, J. S., Krakow, K., & Kleinschmidt, A. (2006). Where the BOLD signal goes when alpha EEG leaves. *NeuroImage*, *31*(4), 1408–1418. https://doi.org/10.1016/j.neuroimage.2006.02.002

Lebel, C., Walker, L., Leemans, A. [A.], Phillips, L., & Beaulieu, C. (2008). Microstructural maturation of the human brain from childhood to adulthood. *NeuroImage*, *40*(3), 1044–1055. https://doi.org/10.1016/j.neuroimage.2007.12.053

Lindsley, D. B. (1939). A Longitudinal Study of the Occipital Alpha Rhythm in Normal Children: Frequency and Amplitude Standards. *The Pedagogical Seminary and Journal of Genetic Psychology*, *55*(1), 197–213. https://doi.org/10.1080/08856559.1939.10533190

Lopes da Silva, F. [F.H], van Lierop, T., Schrijer, C., & van Storm Leeuwen, W. (1973). Organization of thalamic and cortical alpha rhythms: Spectra and coherences. *Electroencephalography and Clinical Neurophysiology*, *35*(6), 627–639. https://doi.org/10.1016/0013-4694(73)90216-2

Lopes da Silva, F. [Fernando] (1991). Neural mechanisms underlying brain waves: from neural membranes to networks. *Electroencephalography and Clinical Neurophysiology*, *79*(2), 81–93. https://doi.org/10.1016/0013-4694(91)90044-5

Luque Laguna, P. A., Combes, A. J. E., Streffer, J., Einstein, S., Timmers, M., Williams, S. C. R., & Dell'Acqua, F. (2020). Reproducibility, reliability and variability of FA and MD in the older healthy population: A test-retest multiparametric analysis. *NeuroImage. Clinical*, *26*, 102168. https://doi.org/10.1016/j.nicl.2020.102168

Marcuse, L. V., Schneider, M., Mortati, K. A., Donnelly, K. M., Arnedo, V., & Grant, A. C. (2008). Quantitative analysis of the EEG posterior-dominant rhythm in healthy adolescents. *Clinical Neurophysiology*, *119*(8), 1778–1781. https://doi.org/10.1016/j.clinph.2008.02.023

Matthis, P., Scheffner, D., Benninger, C. [Chr.], Lipinski, C., & Stolzis, L. (1980). Changes in the background activity of the electroencephalogram according to age. *Electroencephalography and Clinical Neurophysiology*, *49*(5-6), 626–635. https://doi.org/10.1016/0013-4694(80)90403-4

Mazzetti, C., Staudigl, T., Marshall, T. R., Zumer, J. M., Fallon, S. J., & Jensen, O. (2019). Hemispheric Asymmetry of Globus Pallidus Relates to Alpha Modulation in Reward-Related Attentional Tasks. *Journal of Neuroscience*, *39*(46), 9221–9236. https://doi.org/10.1523/JNEUROSCI.0610-19.2019

McIntosh, A. R. [Anthony R.] (2010). The Development of a Noisy Brain. *Archives Italiennes De Biologie*. (148), 223–337. https://doi.org/10.4449/aib.v148i3.1225

Mierau, A., Felsch, M., Hülsdünker, T., Mierau, J., Bullermann, P., Weiß, B., & Strüder, H. K. (2016). The interrelation between sensorimotor abilities, cognitive performance and individual EEG alpha peak frequency in young children. *Clinical Neurophysiology : Official Journal of the International Federation of Clinical Neurophysiology*, *127*(1), 270–276. https://doi.org/10.1016/j.clinph.2015.03.008

Miller, K. J., Sorensen, L. B., Ojemann, J. G., & den Nijs, M. (2009). Power-law scaling in the brain surface electric potential. *PLoS Computational Biology*, *5*(12), e1000609. https://doi.org/10.1371/journal.pcbi.1000609

Mori, S., Crain, B. J., Chacko, V. P., & van Zijl, P. C. M. (1999). Three-dimensional tracking of axonal projections in the brain by magnetic resonance imaging. *Annals of Neurology*, *45*(2), 265–269. https://doi.org/10.1002/1531-8249(199902)45:2<265::AID-ANA21>3.0.CO;2-3

Niedermeyer, E. (1999). The normal EEG of the waking adult. *Electroencephalography: Basic Principles, Clinical Applications, and Related Fields*, *167*, 155–164.

Nyholt, D. R. (2004). A simple correction for multiple testing for single-nucleotide polymorphisms in linkage disequilibrium with each other. *American Journal of Human Genetics*, *74*(4), 765–769. https://doi.org/10.1086/383251

Oldfield, R. C. (1971). The assessment and analysis of handedness: The Edinburgh inventory. *Neuropsychologia*, *9*(1), 97–113. https://doi.org/10.1016/0028-3932(71)90067-4

Pedroni, A., Bahreini, A., & Langer, N. (2019). Automagic: Standardized preprocessing of big EEG data. *NeuroImage*, 460–473. https://doi.org/10.1016/j.neuroimage.2019.06.046

Pion-Tonachini, L., Kreutz-Delgado, K., & Makeig, S. (2019). Iclabel: An automated electroencephalographic independent component classifier, dataset, and website. *NeuroImage*, *198*, 181–197. https://doi.org/10.1016/j.neuroimage.2019.05.026

Posthuma, D., Neale, M. C., Boomsma, D. I., & Geus, E. J. de (2001). Are smarter brains running faster? Heritability of alpha peak frequency, IQ, and their interrelation. *Behavior Genetics*, *31*(6), 567–579. https://doi.org/10.1023/a:1013345411774

Segalowitz, S. J., Santesso, D. L., & Jetha, M. K. (2010). Electrophysiological changes during adolescence: A review. *Brain and Cognition*, *72*(1), 86–100. https://doi.org/10.1016/j.bandc.2009.10.003

Smit, C. M., Wright, M. J., Hansell, N. K., Geffen, G. M., & Martin, N. G. (2006). Genetic variation of individual alpha frequency (IAF) and alpha power in a large adolescent twin sample. *International Journal of Psychophysiology : Official Journal of the International Organization of Psychophysiology*, *61*(2), 235–243. https://doi.org/10.1016/j.ijpsycho.2005.10.004

Smith, S. M. (2002). Fast robust automated brain extraction. *Human Brain Mapping*, *17*(3), 143–155. https://doi.org/10.1002/hbm.10062

Somsen, R. J., van't Klooster, B. J., van der Molen, M. W., van Leeuwen, H. M., & Licht, R. (1997). Growth spurts in brain maturation during middle childhood as indexed by EEG power spectra. *Biological Psychology*, *44*(3), 187–209. https://doi.org/10.1016/S0301-0511(96)05218-0

Surwillo, W. (1961). Frequency of the 'Alpha' Rhythm, Reaction Time and Age. *Nature*, *191*(4790), 823–824. https://doi.org/10.1038/191823a0

Tange, O. (2011). Gnu parallel-the command-line power tool. *The USENIX Magazine*. (36 (1)), 42–47.

Usher, Stemmler, & Olami (1995). Dynamic pattern formation leads to 1/f noise in neural populations. *Physical Review Letters*, *74*(2), 326–329. https://doi.org/10.1103/PhysRevLett.74.326

Vehtari, A., Gelman, A., & Gabry, J. (2017). Practical Bayesian model evaluation using leave-one-out cross-validation and WAIC. *Statistics and Computing*, *27*(5), 1413–1432. https://doi.org/10.1007/s11222-016-9696-4

Veraart, J., Fieremans, E., & Novikov, D. S. (2016). Diffusion MRI noise mapping using random matrix theory. *Magnetic Resonance in Medicine*, *76*(5), 1582–1593. https://doi.org/10.1002/mrm.26059

Veraart, J., Poot, D. H. J., van Hecke, W., Blockx, I., van der Linden, A., Verhoye, M., & Sijbers, J. (2011). More accurate estimation of diffusion tensor parameters using diffusion Kurtosis imaging. *Magnetic Resonance in Medicine*, *65*(1), 138–145. https://doi.org/10.1002/mrm.22603

Veraart, J., Sijbers, J., Sunaert, S., Leemans, A. [Alexander], & Jeurissen, B. (2013). Weighted linear least squares estimation of diffusion MRI parameters: Strengths, limitations, and pitfalls. *NeuroImage*, *81*, 335–346. https://doi.org/10.1016/j.neuroimage.2013.05.028

Voytek, B. [Bradley], & Knight, R. T. [Robert T.] (2015). Dynamic network communication as a unifying neural basis for cognition, development, aging, and disease. *Biological Psychiatry*, *77*(12), 1089–1097. https://doi.org/10.1016/j.biopsych.2015.04.016

Voytek, B. [Bradley], Kramer, M. A., Case, J., Lepage, K. Q., Tempesta, Z. R., Knight, R. T. [Robert T.], & Gazzaley, A. (2015). Age-Related Changes in 1/f Neural Electrophysiological Noise. *Journal of Neuroscience*, *35*(38), 13257–13265. https://doi.org/10.1523/JNEUROSCI.2332-14.2015

Wakana, S., Caprihan, A., Panzenboeck, M. M., Fallon, J. H., Perry, M., Gollub, R. L., . . . Mori, S. (2007). Reproducibility of quantitative tractography methods applied to cerebral white matter. *NeuroImage*, *36*(3), 630–644. https://doi.org/10.1016/j.neuroimage.2007.02.049

Wantanabe, S. (2013). A widely applicable Bayesian information criterion. *Journal of Machine Learning Research*. (14), 867–897. Retrieved from https://www.jmlr.org/papers/volume14/watanabe13a/watanabe13a.pdf

Wassermann, D., Rathi, Y., Bouix, S., Kubicki, M., Kikinis, R., Shenton, M., & Westin, C.-F. (2011). White Matter Bundle Registration and Population Analysis Based on Gaussian Processes. In (pp. 320–332). Springer, Berlin, Heidelberg. https://doi.org/10.1007/978-3-642-22092-0_27

Wechsler, D. (2003). Wechsler intelligence scale for children--Fourth Edition (WISC-IV). *The Psychological Corporation*.

Wechsler, D. (2008). Wechsler Adult Intelligence Scale--Fourth edition (WAIS-IV), Australian and New Zealand Language Adaptation. *NCS Pearson Inc*.

Weiß, R. H. (2011). *Wiener Matrizen-Test 2: Ein Rasch-skaldierter sprachfreier Kurztest zu Erfassung der Intelligenz.*

Welch, P. (1967). The use of fast Fourier transform for the estimation of power spectra: A method based on time averaging over short, modified periodograms. *IEEE Transactions on Audio and Electroacoustics*, *15*(2), 70–73. https://doi.org/10.1109/TAU.1967.1161901

Wen, H., & Liu, Z. (2016). Separating Fractal and Oscillatory Components in the Power Spectrum of Neurophysiological Signal. *Brain Topography*, *29*(1), 13–26. https://doi.org/10.1007/s10548-015-0448-0

Whitford, T. J., Rennie, C. J., Grieve, S. M., Clark, C. R., Gordon, E., & Williams, L. M. (2007). Brain maturation in adolescence: Concurrent changes in neuroanatomy and neurophysiology. *Human Brain Mapping*, *28*(3), 228–237. https://doi.org/10.1002/hbm.20273

Yeatman, J. D., Dougherty, R. F., Myall, N. J., Wandell, B. A., & Feldman, H. M. (2012). Tract profiles of white matter properties: Automating fiber-tract quantification. *PLOS ONE*, *7*(11), e49790. https://doi.org/10.1371/journal.pone.0049790

Yeatman, J. D., Dougherty, R. F., Rykhlevskaia, E., Sherbondy, A. J., Deutsch, G. K., Wandell, B. A., & Ben-Shachar, M. (2011). Anatomical properties of the arcuate fasciculus predict phonological and reading skills in children. *Journal of Cognitive Neuroscience*, *23*(11), 3304–3317. https://doi.org/10.1162/jocn_a_00061

Ystad, M., Hodneland, E., Adolfsdottir, S., Haász, J., Lundervold, A. J., Eichele, T., & Lundervold, A. (2011). Cortico-striatal connectivity and cognition in normal aging: A combined DTI and resting state fMRI study. *NeuroImage*, *55*
